# Exosite inhibition of A Disintegrin And Metalloproteinase with Thrombospondin motif (ADAMTS)-5 by a glycoconjugated arylsulfonamide

**DOI:** 10.1101/2020.06.12.146324

**Authors:** Santamaria Salvatore, Doretta Cuffaro, Elisa Nuti, Lidia Ciccone, Tiziano Tuccinardi, Francesca Liva, Felicia D’Andrea, Rens de Groot, Armando Rossello, Josefin Ahnström

**Affiliations:** Department of Immunology and Inflammation, Imperial College London, Du Cane Road, W12 0NN, London, United Kingdom; Department of Pharmacy, University of Pisa, via Bonanno 6, 56126 Pisa, Italy; Institute of Cardiovascular Science, University College London, 51 Chenies Mews, WC1E 6HX, London, United Kingdom

## Abstract

ADAMTS-5 is a major protease involved in the turnover of proteoglycans such as aggrecan and versican. Its aggrecanase activity has been directly linked to the etiology of osteoarthritis (OA), identifying ADAMTS-5 as a pharmaceutical target for OA treatment. However, most existing ADAMTS-5 inhibitors target its active site and therefore suffer from poor selectivity. Here, using a novel approach, we have designed a new class of sugar-based arylsulfonamide inhibitors, which are selective for ADAMTS-5 through binding to a previously unknown substrate-binding site (exosite). Docking calculations combined with molecular dynamics simulations demonstrated that our lead compound is a cross-domain inhibitor that targets the interface of the metalloproteinase and disintegrin-like domains. Targeted mutagenesis identified disintegrin-like domain residues K532 and K533 as an exosite which is critical for substrate recognition. Furthermore, we show that this exosite acts as major determinant for inhibitor binding and, therefore, can be targeted for development of selective ADAMTS-5 inhibitors.

## Introduction

A Disintegrin And Metalloproteinase with Thrombospondin motif (ADAMTS)-5 (aggrecanase-2) is a metalloproteinase that is important for the proteolytic regulation of the large-aggregating proteoglycans (PGs) aggrecan and versican. These two PGs are responsible for the viscoelastic properties of cartilaginous tissues and large blood vessels, respectively, which are mediated by the glycosaminoglycan (GAG) chains attached to their protein core.^**1,2**^ Dysregulated proteolysis of aggrecan leads to loss of mechanical resilience of cartilage and is therefore a landmark of degenerative pathologies such as osteoarthritis (OA).^**2,3**^ *In vitro*, ADAMTS-5 is the most potent protease against both aggrecan and versican, being 20-30 fold more active than ADAMTS-4 (aggrecanase-1), its closest family member.^**4,5**^ In contrast to *Adamts-4* null mice,^**6**^ *Adamts-5* null mice are protected from cartilage degradation in mechanical and inflammatory OA models.^**7,8**^ Furthermore, inhibitory antibodies against ADAMTS-5 can block aggrecan degradation both in *ex vivo* and *in vivo* in models of OA.^**9,10**^ From a pharmaceutical perspective, ADAMTS-5 is therefore a promising therapeutic target for the treatment of degenerative joint diseases such as OA. However, like many members of the ADAMTS family, ADAMTS-5 is a challenging protein to study due to low expression levels, association with the extracellular matrix, complex post-translational regulation and amenability to (auto)proteolytic degradation.^**4,11**^ It is comprised of a pro-domain (Pro), a metalloproteinase (Mp) domain, a disintegrin-like (Dis) domain, a central thrombospondin-like (Ts) motif, a cysteine-rich (CysR) domain, a spacer (Sp) domain and a C-terminal Ts motif. The Mp domain contains a conserved sequence (<u>H</u>EIG<u>H</u>LLGLS<u>H</u>) where three histidine residues coordinate the zinc ion that is necessary for catalysis. To date, only the crystal structures of the Mp and Dis domains have been solved.^**12,13**^ The lack of structural data for the C-terminal domains has hampered the development of small molecule inhibitors targeting these regions. As a result, all reported ADAMTS-5 synthetic inhibitors act through chelation of the catalytic zinc ion in the Mp domain. The vast majority of current inhibitors contain a zinc-binding group (ZBG), such as hydroxamate or carboxylate.^**14**^ Common drawbacks exhibited by such inhibitors include poor selectivity due to cross-inhibition of other zinc-containing metalloproteinases, high rate of metabolic conversion and poor bioavailability after oral administration. Combined, these are all factors that have limited pre-clinical applications.^**14**^ A strategy to circumvent these drawbacks would be to target specific amino acid residues in the ancillary domains that are necessary for recognition and binding of PGs (i.e. so-called exosites). We have recently identified two such exosites in the ADAMTS-5 Sp domain (residues 739-744 and 837-844).^**5**^ These regions are essential for proteolysis of both aggrecan and versican and lie in loops which are hypervariable within the ADAMTS family,^**5**^ thus offering a target for the development of selective inhibitors. However, exposed loops on metalloproteinases can be more easily targeted by antibodies than by small molecules,^**15**^ while small molecule inhibitors tend to bind hydrophobic crevices.^**16**^ GAGs represent a potential third avenue to exosite inhibition. For example, heparin, a naturally occurring GAG, inhibits the proteoglycanase activity of ADAMTS-5 by binding to the Sp and CysR domains.^**4,11,17**^ However, due to its anticoagulant properties and associated side effects, such as bleeding^**18**^ and thrombocytopenia,^**19**^ heparin itself is not suitable as a therapeutic inhibitor. In heparin, the most common disaccharide unit is composed of a 2-*O*-sulfated iduronic acid and 6-*O*-sulfated, *N*-sulfated β-*N*-acetyl-D-glucosamine (GlcNAc), which we here have exploited for the development of an ADAMTS-5 inhibitor. For this purpose, we probed ADAMTS-5 with a small library of GAG-mimetic molecules, containing the GlcNAc moiety (**Figure 1A**). Starting from our previously described zinc-chelating inhibitors, with no selectivity for ADAMTS-5 *versus* ADAMTS-4,^**20**^ we successfully identified an ADAMTS-5-selective glycoconjugate, devoid of a ZBG. Furthermore, we show that the GlcNAc moiety of the inhibitor binds to a cluster of positively charged residues in the ADAMTS-5 Dis domain. Through mutagenesis and biochemical characterization, we also show that these residues compose a novel exosite which is amenable to selective inhibition.

**Figure 1.**
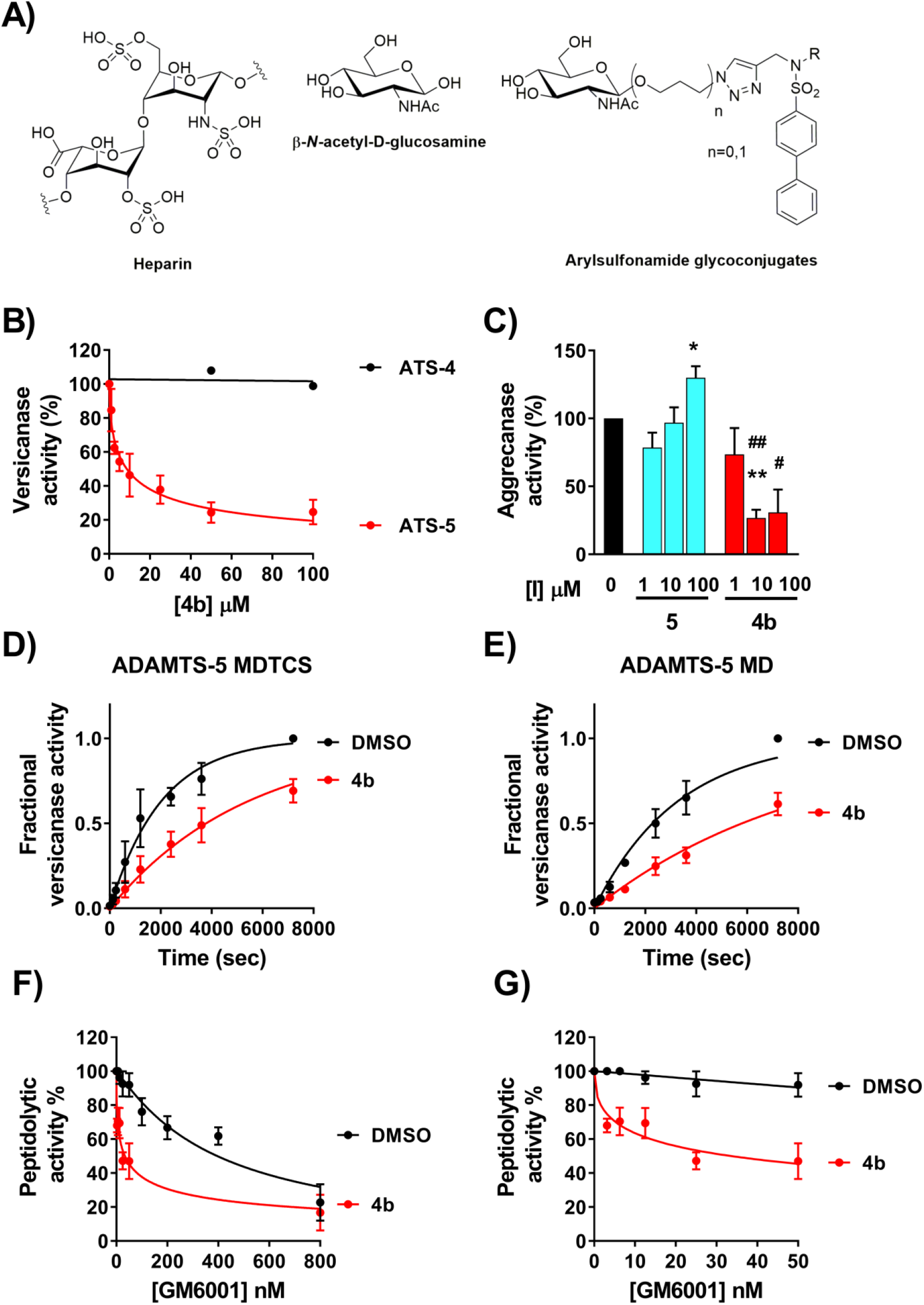
Characterization of the inhibitory activity of compound 4b. **A)** Structural comparison between heparin and the glycoconjugates analyzed for ADAMTS-5 inhibition in this study. R represents either a ZBG or an aliphatic moiety. **B)** Inhibition of ADAMTS-5 versicanase activity by 4b. ADAMTS-4 (5.5 nM) and −5 (0.4 nM) were incubated either with compound 4b or DMSO for 2 h at 37°C before addition of V1-5GAG (50 nM). At each time point, reactions were stopped by addition of EDTA and ADAMTS-5 generated versican fragments (versikine) quantified by sandwich ELISA. The relative versicanase activity is presented and 100% activity corresponds to that in the presence of DMSO alone. **C)** Analysis of inhibition of ADAMTS-5 aggrecanase activity by 4b. Compounds 4b and 5 were incubated with ADAMTS-5 (1 nM) for 2 h at 37°C before addition of aggrecan (20 μg). Following SDS-PAGE and immunoblot, fragments cleaved at the Glu392↓Ala393 bond were detected by a monoclonal neoepitope antibody recognizing the new C-terminal fragment (anti-ARGSV) and analyzed by densitometric analysis. Data are presented as mean ± SEM (n=4). *p<0.005 and **p<0.001, compared to DMSO controls; #p<0.005 and ##p<0.001, compared to the same concentration of compound 5 (Mann-Whitney test). **D, E)** Compound 4b inhibits the ADAMTS-5 activity by binding to its Mp/Dis domains. Compound 4b (100 μM) or DMSO was incubated either with ADAMTS-5 full-length (FL) (0.2 nM, C) or ADAMTS-5 MD (26 nM, D) for 2h at 37°C before addition of V1-5GAG (50 nM). At each time point, reactions were stopped by addition of EDTA and versikine fragments quantified by sandwich ELISA as reported in the Method section. In the DMSO control, complete proteolysis was achieved after 7200 seconds. **F,G)** Compound 4b does not bind to the active site zinc. ADAMTS-5 MDTCS was incubated with active-site inhibitor GM6001 (0-800 nM) either in the presence of DMSO or compound 4b (10 μM) for 2 h at 37°C before addition of QF-peptide. The synergistic effect exerted by compound 4b is more evident at low concentrations of GM6001 (panel G). The relative peptidolytic activity is presented and 100% activity corresponds to that in the presence of DMSO alone. Data are presented as mean ± SEM (n=3).

## Results

### Development of an ADAMTS-5 exosite inhibitor

The primary aim of this study was to develop a selective ADAMTS-5 exosite inhibitor. Exosites are located away from the active site and therefore most of them do not bind small synthetic peptides that are used in the standard Quenched-Fluorescent (QF)-peptide cleavage assays.^**9**^ Therefore, to accurately measure inhibition by *exosite* inhibitors, we used our novel ELISA-based assay that employs V1-5GAG, a truncated versican V1 variant, as a substrate. This contains all the binding sites for efficient cleavage by ADAMTS-5 and allows precise kinetic quantification of inhibition.^**5, 21**^

To develop ADAMTS-5 exosite inhibitors, we started with heparin mimetics (GlcNAc based) (**Figure 1A**) fused to an arylsulfonamide scaffold containing a weak ZBG (carboxylic acid).^**20**^ In the second stage, we removed the ZBG to increase selectivity. We first performed a small structure-activity relationship (SAR) study with these glycoconjugated arylsulfonamides having either the GlcNac directly attached to the arylsulfonamido group or via a *n*-propyloxy linker (**Table 1**). Our previously described inhibitors, compounds **1** and **2**,^**20**^ inhibited ADAMTS-5 and ADAMTS-4 to a similar level, with half maximal inhibitory concentration (IC_50_) values in the micromolar range (**Table 1**). Replacement of the carboxylate with a benzyl ester completely abolished inhibitory activity in both series (compounds **3a** and **4a**), likely due to steric hindrance. Importantly, replacement of the carboxylic acid ZBG with a *sec*-butyl group to generate compound **4b** improved ADAMTS-5 inhibition against V1-5GAG approximately 5-fold and, at the same time, abolished ADAMTS-4 inhibition (**Figure 1B** and **Table 1**). This is a remarkable increase in selectivity, in light of the homology and the functional overlap between these two proteases. The presence of an *n*-propyloxy link between the sugar and the arylsulfonamide was essential for inhibition of ADAMTS-5 versicanase activity as shown by the ~13-fold reduction in inhibition when the sugar was directly linked to the sulfonamide scaffold as in compound **3b** (**Table 1**).

**Table 1.**
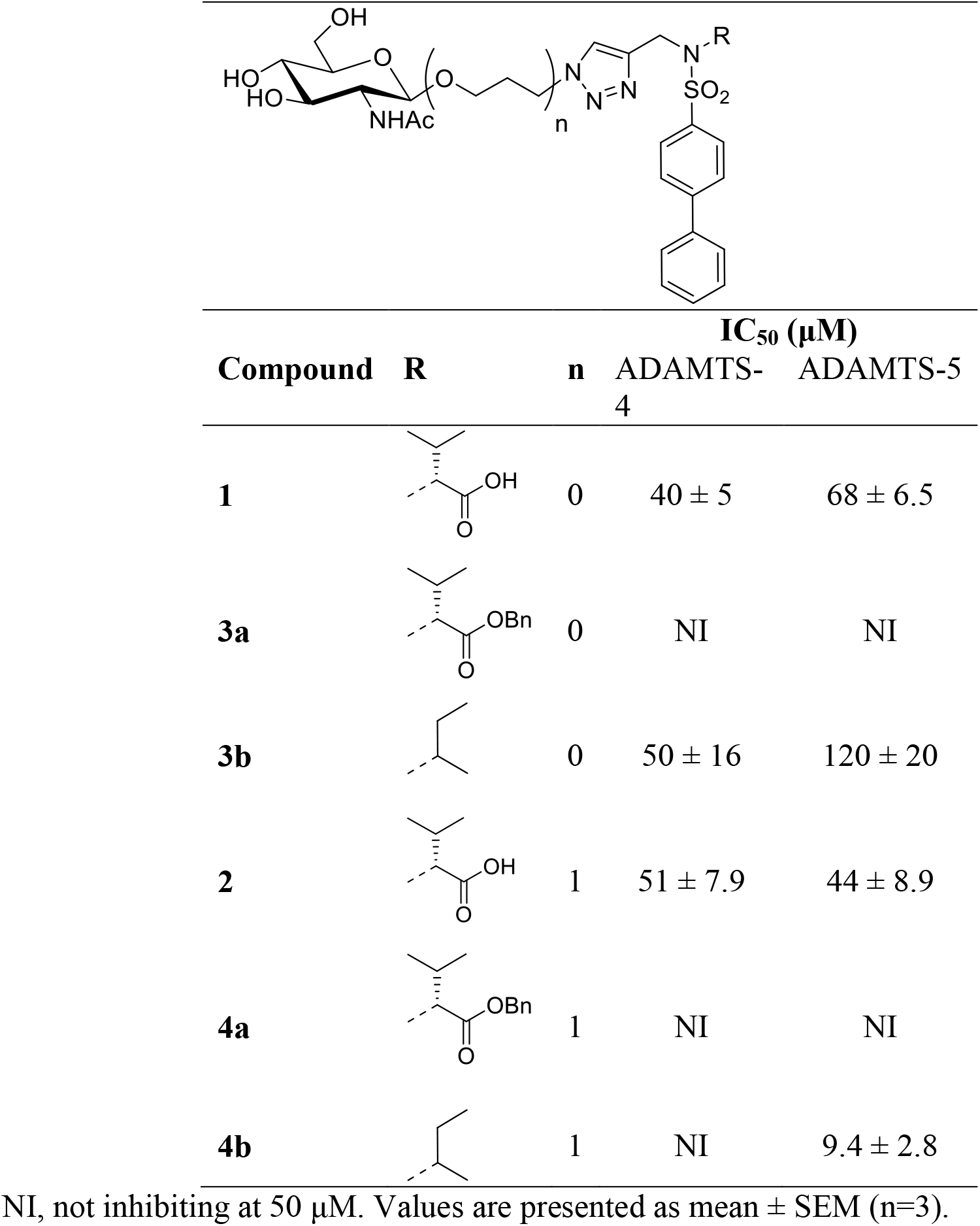
Inhibitory activities of glycoconjugates 1, 2, 3a, 3b, 4a and 4b against ADAMTS-4 and -5.

|  |  |  |  |  |
| --- | --- | --- | --- | --- |
| Compound | R | n | IC<sub>50</sub> (μM) |  |
|  |  |  | ADAMTS-4 | ADAMTS-5 |
| **1** |  | 0 | 40 ± 5 | 68 ± 6.5 |
| **3a** |  | 0 | NI | NI |
| **3b** |  | 0 | 50 ± 16 | 120 ± 20 |
| **2** |  | 1 | 51 ± 7.9 | 44 ± 8.9 |
| **4a** |  | 1 | NI | NI |
| **4b** |  | 1 | NI | 9.4 ± 2.8 |

NI, not inhibiting at 50 μM. Values are presented as mean ± SEM (n=3).

Interestingly, when tested in a QF-peptide cleavage assay (**Table 2**), **4b** did not inhibit ADAMTS-5 peptidolytic activity, suggesting that this compound may target an exosite. To understand the binding mode of the inhibitor, we performed systematic modifications of **4b** (**Table 2**). Replacement of the sugar moiety with a benzoyl group (compound **5**) abolished inhibitory activity, suggesting a direct interaction of the GlcNAc with ADAMTS-5. The introduction of a benzyloxyphenyl in the arylsulfonamide moiety was reported to greatly improve the inhibitory activity of arylsulfonamide hydroxamates against ADAMTS-5.^**22**^ Compounds containing this modification (**4c** and **4d**) showed a severe reduction in inhibitory activity which was also associated with a decreased selectivity over ADAMTS-4 (**Table 2**), suggesting a different mode of interaction for the glycoconjugates.

**Table 2.**
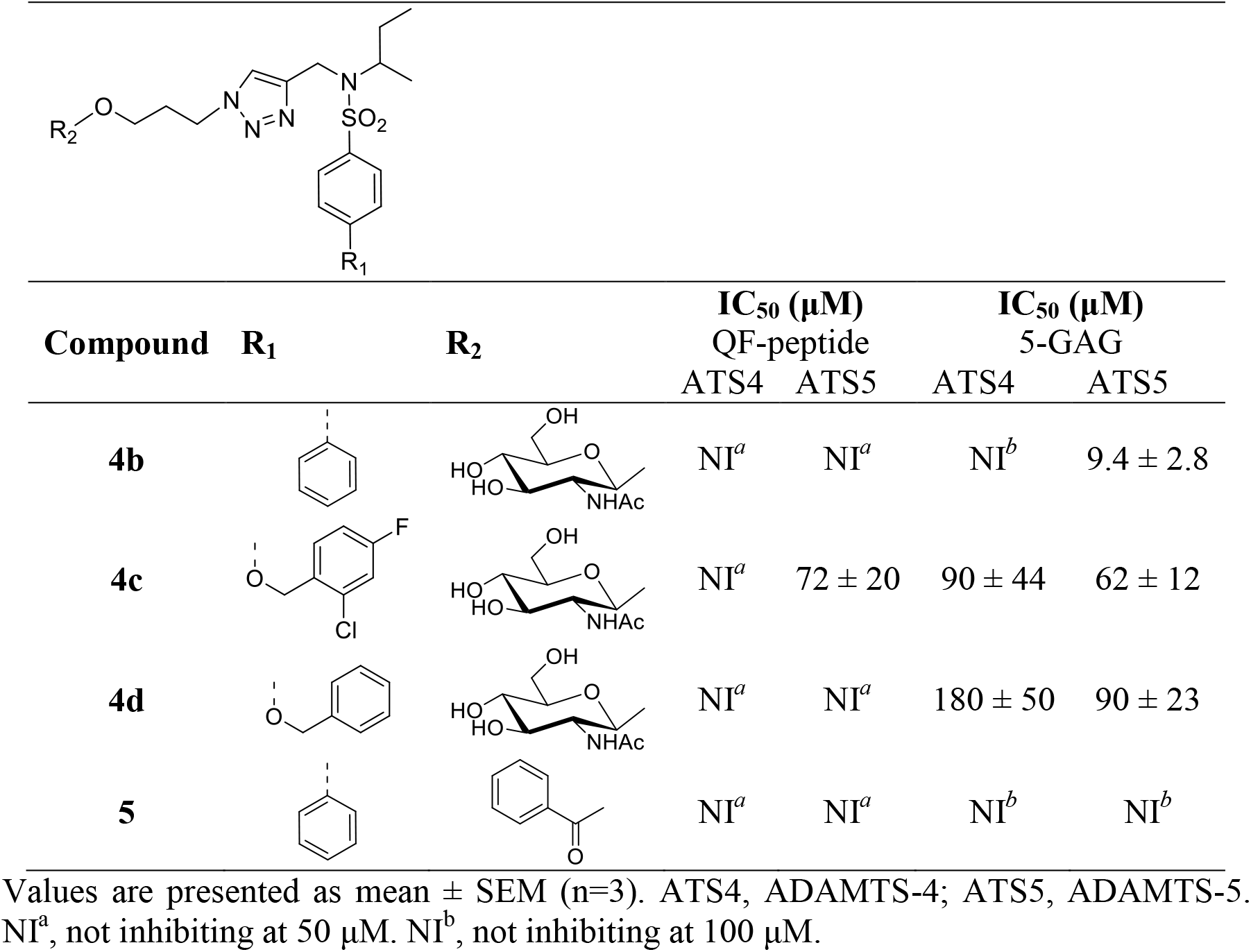
Inhibitory activities of 4b and its derivatives, 4c,d and 5, against ADAMTS-4 and -5.

|  |  |  |  |  |  |  |
| --- | --- | --- | --- | --- | --- | --- |
| Compound | R<sub>1</sub> | R<sub>2</sub> | IC<sub>50</sub> (μM)<br>QF-peptide |  | IC<sub>50</sub> (μM)<br>5-GAG |  |
|  |  |  | ATS4 | ATS5 | ATS4 | ATS5 |
| **4b** |  |  | NI<sup>a</sup> | NI<sup>a</sup> | NI<sup>b</sup> | 9.4 ± 2.8 |
| **4c** |  |  | NI<sup>a</sup> | 72 ± 20 | 90 ± 44 | 62 ± 12 |
| **4d** |  |  | NI<sup>a</sup> | NI<sup>a</sup> | 180 ± 50 | 90 ± 23 |
| **5** |  |  | NI<sup>a</sup> | NI<sup>a</sup> | NI<sup>b</sup> | NI<sup>b</sup> |

Values are presented as mean ± SEM (n=3). ATS4, ADAMTS-4; ATS5, ADAMTS-5. NI<sup>a</sup>, not inhibiting at 50 μM. NI<sup>b</sup>, not inhibiting at 100 μM.

Importantly, the inhibition of ADAMTS-5 by **4b** was confirmed using aggrecan as a substrate. Compound **4b** effectively inhibited aggrecan proteolysis at Glu392↓Ala393 (**Figure 1C and Figure 1 -Figure Supplement 1**), the cleavage site which is most detrimental for cartilage integrity.^**23**^ Approximately 80% ADAMTS-5 inhibition was observed at 10 μM. In contrast, the analogue compound devoid of the sugar moiety, **5**, had no effect upon aggrecan cleavage (**Figure 1C**). These results confirmed that **4b** is suitable for further development as a potential disease-modifying OA agent.

To further probe the interaction of **4b** with ADAMTS-5, we compared its inhibitory activity against ADAMTS-5 MDTCS (lacking the last TS motif but retaining full versicanase activity),^**4**^ and its variant consisting only of the Mp and Dis domains (ADAMTS-5 MD), the minimal variant endowed with detectable proteoglycanase activity (**Figure 1D and 1E**).^**11**^ As previously reported,^**5**^ deletion of the C-terminal ancillary domains reduced ADAMTS-5 versicanase activity by approximately 730-fold (*k*_cat_/*K*_m_: (34.3 ± 0.43) × 10^5^ M^−1^ s^−1^ for ADAMTS-5 MDTCS *versus* (0.14 ± 0.03) × 10^5^ M^−1^ s^−1^ for ADAMTS-5 MD, p<0.001). In the presence of 100 μM of compound **4b,** ADAMTS-5 MDTCS and MD cleaved V1-5GAG with a *k*_cat_/*K*_m_ values of (10.0 ± 0.30) × 10^5^ M^−1^ s^−1^ and (0.047 ± 0.010) × 10^5^ M^−1^ s^−1^, respectively. This corresponds to a 71% and 66% reduction in activity for the MDTCS and MD variants, respectively (p<0.001 for both compared to their controls containing equal amounts v/v of dimethyl sulfoxide, DMSO). The similar extent of inhibition exerted by compound **4b** on the two variants confirms that this inhibitor binds to the Mp/Dis domains.

### Binding mode of compound 4b

Since compound **4b** is devoid of an obvious ZBG and does not have any effect on the ADAMTS-5 peptidolytic activity while still inhibiting its proteoglycanase activity (**Table 2**), we hypothesized that it does not interact with the ADAMTS-5 catalytic zinc. To test this, we performed QF-peptide cleavage assays at increasing concentrations of GM6001 (Ilomastat, 0-800 nM), a known zinc-binder and broad-spectrum metalloproteinase inhibitor,^**22, 24**^ in the presence or absence of 10 μM **4b** (**Figure 1F and 1G**). The inhibitory activity of GM6001 was quantified as an IC_50_ in the presence (IC_50_^+^) and absence (IC_50_^−^) of **4b**. If the IC_50_ of GM6001 was unchanged in the presence of **4b** (IC_50_^+^/IC_50_^−^ ~1) no synergism between the molecules would occur, whereas if the IC_50_ was reduced (IC_50_^+^/IC_50_^−^ <1) or increased (IC_50_^+^/IC_50_^−^ >1), the interaction would be synergistic or antagonistic, respectively. GM6001 showed a large increase in inhibition in the presence of **4b** (IC_50_^+^=16 ± 5 nM/IC_50_^−^ = 370 ± 90 nM; IC_50_^+^/IC_50_^−^ =0.043, p<0.05), and therefore acted as a synergistic inhibitor together with **4b**. This confirms that **4b** does not bind the catalytic zinc but binds close enough to the active site to have a synergistic effect upon GM6001 binding. This is also in agreement with the results using ADAMTS-5 MD (**Figure 1E**) which suggested that the interaction site for **4b** is contained within these two domains.

To investigate how compound **4b** interacts with ADAMTS-5, docking calculations combined with molecular dynamics (MD) simulations were carried out. The compound was docked into the catalytic site of ADAMTS-5 (PDB code: 2RJQ) complexed with GM6001 using AUTODOCK 4.2 software. Two hundred docking poses were generated and then clustered by applying a root-mean-square deviation (RMSD) of 6.0 Å. The cluster analysis suggested three possible dispositions for **4b** (**Figure 2 −Figure supplement 1**). Therefore, a representative docking pose belonging to each of the three cluster of poses (C1-C3) was subjected to a 103 ns MD simulation with explicit water molecules, followed by analysis of the three binding modes through the molecular mechanics and Poisson Boltzmann surface area (MM-PBSA) method. The results obtained from these analyses suggested the MD-refined C1 pose as the most reliable binding disposition of **4b** within ADAMTS-5. As shown in **Figure 2A**, the sugar portion of the molecule interacts with the Dis domain of ADAMTS-5, whereas the remaining parts the molecule interacts with the Mp domain. **Figure 2B** shows the main interactions of **4b** with ADAMTS-5. The biphenyl fragment is water-exposed and shows lipophilic interactions with H374. One of the two oxygen atoms of the sulfonamide moiety forms an hydrogen (H)-bond with the hydroxyl group of S375 whereas the triazole ring shows lipophilic interactions with the indole ring of GM6001. Thus, the results obtained from the docking experiments provide a mechanistic rationale for the synergistic effect observed in the QF-peptide cleavage assay. Finally, a particularly important finding was that the GlcNAc showed H-bond interactions with ADAMTS-5 Dis domain residues K532 and K533. The length of the linker appears to be critical in positioning the GlcNAc group at the right distance from the Dis domain, as also shown by the reduced inhibitory potency and selectivity exhibited by compound **3b,** where the sugar is directly linked to the sulfonamide group (**Table 1**).

**Figure 2.**
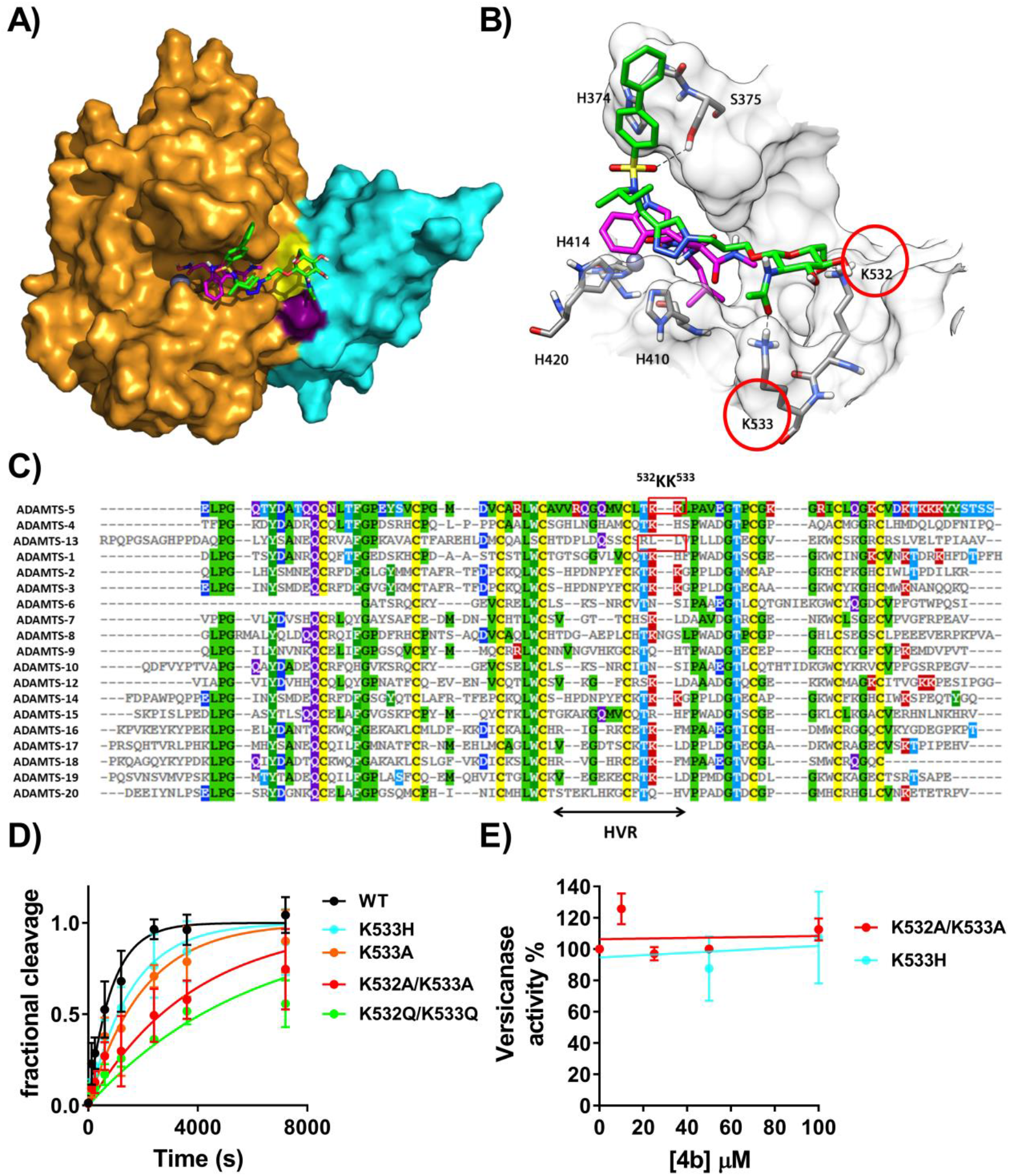
Compound 4b binds to an exosite in the ADAMTS-5 Dis domain. **A, B**) Minimized average structure of compound 4b bound to the ADAMTS-5-GM6001 complex, derived from the last 100 ns of MD simulation. (A) Protein surface. 4b is shown in green, GM6001 is in pink, ADAMTS-5 Mp domain in bright orange, the Dis domain in light blue, the active site zinc as a grey sphere. Exosite residues K532 and K533 are colored in yellow and purple, respectively. (B) Protein residues directly interacting with compound 4b. H-bonds are represented as black dashed lines. (**C**) Amino acid sequence alignment of the Dis domain of all ADAMTS family members. Known exosites are boxed and the poorly conserved region (HVR, hypervariable region) is indicated. Conserved residues are indicated by the same color code. **D**) Time course experiments for cleavage of 50 nM V1-5GAG by ADAMTS-5 MDTCS variants. Each enzyme (0.4 nM) was incubated with 50 nM substrate. At the indicated time points, an aliquot was taken, stopped with EDTA and cleavage products measured by sandwich ELISA as described in the Experimental procedures. The solid lines represent a nonlinear regression fit of the data. The data are presented as average ± SEM; n=3-4. **E**) Inhibition of ADAMTS-5 MDTCS variants by compound 4b. ADAMTS-5 K533H and K532A/K533A (0.82 nM) were incubated either with compound 4b or DMSO for 2h at 37°C before addition of V1-5GAG (50 nM). At each time point, reactions were stopped by addition of EDTA and cleavage fragments quantified by sandwich ELISA. The relative versicanase activity is presented and 100% activity corresponds to that in the presence of DMSO alone. The data are presented as average ± SEM; n=3.

### Identification of an exosite in the ADAMTS-5 Dis domain

Our experimental and *in silico* results both suggested that **4b** functions as a cross-domain exosite inhibitor. They also suggested that residues K532 and K533 compose an undescribed exosite in the ADAMTS-5 Dis domain. To further investigate the importance of K532 and K533 as an exosite for versican cleavage and inhibition by compound **4b**, we substituted both residues either to alanine (variant K532A/K533A) or glutamine (variant K532Q/K533Q) in ADAMTS-5 MDTCS. An amino acid sequence alignment of the Dis domains showed that ADAMTS-4 retains a lysine residue at position 532 but presents a histidine residue at position 533 (**Figure 2C**). Since compound **4b** is selective for ADAMTS-5 over ADAMTS-4, we also substituted K533 to histidine (K533H), to determine how important this residue is for inhibitor selectivity. ADAMTS-5 K533A was generated to assess the overall importance of K533 for substrate recognition and protein stability. All variants were transiently expressed in HEK293T cells. Western blot analysis of conditioned media demonstrated that all mutants were expressed and secreted at similar levels to that of wild-type ADAMTS-5 (**Figure 2 -Figure Supplement 2A**). Following purification (**Figure 2 -Figure Supplement 2B**), these variants were tested in the versican digestion assay (**Figure 2D** and **Table 3**). Interestingly, replacement of the two contiguous lysine residues resulted in a significant reduction in versicanase activity, suggesting that these residues are necessary for recognition and proteolysis of PGs. Critically, compound **4b** was unable to inhibit the versicanase activity of K532A/K533A (**Figure 2E**), thus confirming our *in silico* data. Replacement of K533 either to alanine (to abolish the positive charge on side chain) or histidine (as in ADAMTS-4) resulted only in a modest reduction in versicanase activity. Compound **4b** did not show inhibitory activity against the K533H variant (**Figure 2E**), thus demonstrating that binding of the inhibitor to Lys533 in ADAMTS-5 creates the desired selectivity against ADAMTS-4.

**Table 3.** Kinetic parameters for proteolysis of V1 5-GAG by ADAMTS-5 variants. Values were determined by time course experiments at 50 nM V1-5GAG substrate concentration.

| ADAMTS-5<br>variant | $k_{cat}/K_m^a$<br>$10^5 \text{ M}^{-1} \text{ s}^{-1}$ | Fold<br>reduction |
| --- | --- | --- |
| MDTCS | $32.3 \pm 7.64$ | - |
| K532A/K533A | $6.75 \pm 2.22^*$ | 4.8 |
| K532Q/K533Q | $4.01 \pm 0.49^*$ | 8.0 |
| K533A | $12.5 \pm 2.75^*$ | 2.6 |
| K533H | $15.6 \pm 4.17$ | 2.1 |

Results are expressed as mean ± SEM. \*p<0.05, compared to ADAMTS-5 MDTCS according to Mann-Whitney tests (n=3).

## Discussion

Cumulative evidence from the last 20 years has recognized ADAMTS-5 as a target for OA. Currently, three clinical trials (ID: NCT03595618, NCT03583346 and NCT03224702) are under way to assess the efficacy of ADAMTS-5 inhibitors as OA therapeutics. These involve a small molecule containing a ZBG and a monoclonal antibody.^**25**^ However, many inhibitors have failed at the preclinical stage,^**25**^ a major reason being the lack of adequate selectivity. To create a break-through, we aimed to alter the principle behind the inhibition from active site inhibition to exosite inhibition. This was made possible by our kinetic assay, using versican instead of peptides as a substrate.^**9**^ Our inhibitor design strategy involved conjugating GlcNAc to a ZBG-arylsulfonamide scaffold, followed by removal of the ZBG. This strategy diverges from the traditional approach where ZBGs are attached to an S1’ binding moiety.^**26**^ We confirmed that our lead compound, **4b** does not bind to the active site zinc, as shown by the lack of inhibition of QF-peptide cleavage, as well as its synergism with the zinc-chelating broad-spectrum metalloproteinase inhibitor GM6001. Replacement of the GlcNAc with a benzoyl group (**Table 2**) abolished inhibitory activity, suggesting a direct interaction of the GlcNAc with ADAMTS-5. To our knowledge, a similar role of an amino sugar moiety has not been reported previously for metzincin inhibitors. Instead, the addition of a carbohydrate group has been envisaged as a way to increase the hydrophilicity of metalloproteinase inhibitors and thus enhance their oral availability, without affecting their inhibitory activity. ^**20, 27–30**^ Using a combination of kinetic and *in silico* studies, we demonstrated that **4b** is an exosite cross-domain inhibitor, acting by an unprecedented mechanism where the S1’ pocket is occupied by the arylsulfonamide portion, whereas the sugar moiety interacts with the exosite in the Dis domain. Overall, the binding of **4b** may have two consequences: 1) to occlude access of PGs to the active site; 2) “freezing” the flexibility between the Mp and Dis domains in a way similar to the inhibitory antibody described by Larkin et al.^**10**^

In contrast to what their names suggest, the Dis domains of ADAMTS family proteases do not share homology with disintegrins, a family of proteins from viper venoms. Instead, they are structurally similar to a region in the CysR domains of P-III snake venom metalloproteinases which are involved in binding to platelet integrin receptors.^**12, 13, 31–33**^ In ADAMTS-5, the Dis domain lacks any integrin binding sequence and does also not interact with integrins.^**13**^ It has a unique fold of two α-helices, two β-sheets, and several loops throughout the domain and is connected to the Mp domain by a flexible linker that is 9 amino acids long.^**13**^ The Dis domain lies on the prime side of the active site, where it shields the S1’ pocket and, to a lesser extent, the S3’ pocket from solvent.^**13**^ Previous studies have shown that the isolated ADAMTS-5 Mp domain alone is unable to cleave PGs. ^**11, 21**^ Its proteoglycanase activity is partially restored when the Dis domain is added (ADAMTS-5 MD), although addition of further C-terminal ancillary domain is necessary for full proteoglycanase activity. ^**4, 5**^ The same lack of proteolytic activity of the isolated Mp domain in the absence of the Dis domain, has been observed for several other ADAMTS family members, such as ADAMTS-1,^**34**^ −4,^**35**^ −9,^**36**^ and −13.^**37, 38**^ Together, these results suggest that the Dis domain may be involved in substrate recognition for all five mentioned ADAMTS members.

The GlcNAc group in inhibitor **4b** interacts with residues K532 and K533 in the Dis domain where they constitute a previously undescribed exosite. These two residues lie adjacent to the active site cleft, at a distance of 17-18 Å from the active site zinc, in a region poorly conserved (hypervariable) amongst the ADAMTS family of proteases (**Figure 2C**). The ADAMTS-5 Dis exosite partially overlaps with an exosite in ADAMTS-13 (residues R349, L350G and V352G).^**39**^ However, the distance between the Dis exosite and the zinc is higher in ADAMTS-13 (~26Å), possibly to adapt to its specific substrate, von Willebrand factor.^**39**^ Moreover, the ADAMTS-5 exosite is more positively-charged, suggesting that the interaction with PGs involves more electrostatic than hydrophobic interactions. This hypothesis is supported by the presence of a lysine-rich sequence (^739^NKKSKG^744^) in the ADAMTS-5 Sp domain which has been suggested to bind heparin^**17**^ and PGs,^**5**^ suggesting similarities between the amino acid composition of the exosites present in the Sp and Dis domains. Semisynthetic polysaccharides such as pentosan polysulfate have also been shown to bind to ADAMTS-5 Mp/Dis through electrostatic interactions.^**17**^ Kinetic analysis revealed a 5-8 -fold reduction in catalytic efficiency when both K532 and K533 where mutated, thus showing their importance for efficient substrate cleavage. However, this effect is modest compared with that observed after substitutions of the β1-β2 and β9-β10 loops in the Sp domain (residues 739-744 and 837-844, respectively), which caused a 30-40 fold reduction in versicanase activity, compared to wild-type ADAMTS-5.^**5**^ It is therefore likely that the contact between the Dis exosite and PGs follows an initial binding of the PGs to the ADAMTS-5 Sp domain. ^**5**^ Other studies have also suggested the functional importance of the Dis domain. For example, the endogenous inhibitor of ADAMTS-5, Tissue Inhibitor of Metalloproteinase (TIMP)-3, interacts with the Dis domain^**9, 40**^ and monoclonal antibodies, inhibiting ADAMTS-5 function, have also been reported that target this domain.^**9, 10**^ However, to date compound **4b** is the only example of a small molecule binding to the Dis domain. Further optimization of this cross-domain sugar-based exosite inhibitor could lead to the development of a novel class of OA therapeutics with increased selectivity and bioavailability.

## Materials and Methods

### Protein expression and purification

The constructs coding for human ADAMTS-4 and −5 with a C-terminal FLAG tag (DYKDDDDK) in pEGFP-N1 vector have been described previously.^**5**^ ADAMTS-5 variants were generated using site-directed mutagenesis and confirmed through sequencing. The versican V1-5GAG plasmid, comprising amino acids 21-694 of V1 with C-terminal C-myc/6x His tag has been described previously. ^**5,21**^ Expression and purification of ADAMTS-5 variants and V1-5GAG was performed as previously reported. ^**5**^ DNA and protein concentrations were measured using a NanoDrop ND-2000 UV-visible spectrophotometer (Thermo Fisher Scientific, Nottingham, UK).

### Inhibition assays

All enzyme assays were conducted in TNC-B buffer (50 mM Tris-HCl, pH 7.5, 150 mM NaCl, 10 mM CaCl_2_, 0.05% Brij-35 and 0.02% NaN_3_) at 37°C. The inhibitor stock solutions in DMSO (10 mM) were diluted at different concentrations and incubated with ADAMTS-4 or ADAMTS-5 for 2 h at 37°C. Percent of inhibition was calculated from control reactions containing only DMSO. IC_50_ values were determined using the formula: *v*_i_/*v*_0_ =1/(1 + [I]/IC_50_) where *v*_i_ is the initial velocity of substrate cleavage in the presence of the inhibitor at concentration [I] and *v*_0_ is the initial velocity in the presence of an equal concentration (v/v) of DMSO.

### Versican digestion assays

ADAMTS-5 MDTCS (final concentration 0.4 nM), ADAMTS-5 MD (26 nM) or ADAMTS-4 (5.5 nM) were incubated with different concentrations of inhibitors or DMSO for 2 h at 37°C in TNC-B buffer before addition of V1-5GAG (50 nM). At different time points (0-20 min), sub-samples were removed and reactions were stopped with EDTA. Maxisorp plates (VWR, Lutterworth, UK) were coated with 5 μg/mL anti-DPEEAE neoepitope antibody (Cat n. PA1-1748A, Life Technologies, Paisley, UK) in carbonate buffer pH 9.6 (16 h, 4°C). This neoepitope antibody specifically recognizes the N-terminal versican fragment versikine, generated when ADAMTS-5 cleaves versican at the Glu441↓442Ala bond. Washing steps were performed in triplicate with 300 μL phosphate buffered-saline (PBS) containing 0.1% Tween-20 between each step. Plates were blocked with 3% bovine serum albumin (BSA)/PBS for 2 h, at RT. The samples from the digestion experiments were diluted in 3% BSA/PBS and added to the plate (100 μL, 2h, RT). Bound versikine fragments were detected using anti-G1 monoclonal antibody (Cat n. ab171887, Abcam, Cambridge, UK) (3 μg/mL in 0.5% BSA/PBS, 1.5 h, RT), followed by horseradish peroxidase (HRP)-conjugated anti-mouse antibodies (Cat. N. P044701-2, Agilent Technologies LTD, Cheadle, UK) (2.4μg/mL, 1h, RT). The assay was developed by addition of o-phenylenediamine dihydrochloride (OPD, Cat n. 34006, Sigma Aldrich, Gillingham, UK) for 10 minutes and reactions were stopped with 2 M H_2_SO_4_. The absorbance was read at 492 nm using a BioTeK Epoch (BioTek, Swindon, UK) plate reader. For each dilution, the amount of neoepitope generated was derived from a standard curve (0-1.56 nM) of V1-5GAG completely digested with ADAMTS-5. Initial velocities were calculated from the concentration of versikine generated as a function of reaction time and IC_50_ values were determined.

For inhibition studies, initial rates of proteolysis (<20% cleavage) were analyzed between 0 and 20 minutes. For determination of the specificity constants (*k*_cat_/*K*_m_), digestion reactions were allowed to occur to completion (0-2 h). Data were analyzed as previously described.^**41**^

### Aggrecan digestion assays

ADAMTS-5 (5 nM) was incubated with inhibitors or DMSO for 2 h at 37°C in TNC-B buffer. Aggrecan from bovine articular cartilage (270 nM) (Cat. n.: A1960 Sigma Aldrich, numbering according to Uniprot accession number: P13608) was added. After 2 h digestion at 37°C, the reactions were stopped with EDTA buffer and the samples incubated with 0.1U/mL of chondroitinase ABC (AMS Biotechnology, Abingdon, UK) and keratanase (endo-beta galactosidase, Cat. n.: G6920, Sigma Aldrich) in deglycosylation buffer (50mM sodium acetate, 25 mM Tris HCl pH 8.0) for 16 h at 37°C to remove GAG chains. Samples were analyzed by SDS-PAGE under reducing conditions (5% β-mercaptoethanol) on 4-12% Bis-Tris NuPage Gels (Thermo Fisher) and cleavage products were detected using mouse monoclonal BC-3 antibody which detects aggrecan cleavage at the Glu392↓Ala393 bond (Cat n.: MA316888, Life Technologies). Immobilon Chemiluminescent HRP substrate (Cat. n. IMGDV002, Merck Millipore, Watford, UK) was used for detection. Bands were detected with a Chemidoc Touch Imaging system (Bio-Rad Laboratories Ltd, Hemel Hempstead, UK) and intensities were measured using Image lab software version 5.2.1.

### QF peptide cleavage assays

QF peptide cleavage assays were conducted in 96-well black microtiter plates (Scientific Laboratories Supplies Ltd, Wilford, UK) using a Fluostar Omega microplate reader (BMG Labtech, Aylesbury, UK). The activity of ADAMTS-4 and −5 was monitored for 2 h using the fluorescent peptide substrates fluorescein 5(6)-carbonyl -Ala-Glu-Leu-Asn-Gly-Arg-Pro-Ile-Ser-Ile-Ala-Lys-N,N,N0,N0-tetramethyl-6-carboxyrhodamine-NH_2_ (FAM-AE~LQGRPISIAK-TAMRA, ADAMTS-4) and fluorescein 5(6)-carbonyl-Thr-Glu-Ser-Glu~Ser-Arg-Gly-Ala-Ile-Tyr-Lys-Lys-N,N,N0,N0-tetramethyl-6-carboxyrhodamine-NH_2_ (FAM-TESE ~SRGAIYKK-TAMRA, ADAMTS-5) (custom-synthesized by Bachem, Bubendorf, Switzerland) with an excitation wavelength of 485 nm and an emission wavelength of 538 nm. Final substrate concentrations were 1 μM and 40 μM for ADAMTS-4 and −5, respectively. Fluorescence was expressed in relative fluorescence units (RFU) and normalized against a blank containing only buffer and substrate. Emission spectra were recorded at several concentrations of each compound to exclude auto-fluorescence and maximal inhibitor concentrations were selected accordingly.

### Dual inhibition studies

Dual inhibitor experiments were performed essentially as before.^**42**^ Briefly, GM6001 (0-800 nM) was incubated with ADAMTS-5 in the presence and absence of a fixed concentration (10 μM) of compound **4b** and QF-peptide cleavage assays were run as above. IC_50_ values were determined in the absence (IC_50_^−^) and in the presence of **4b** (IC_50_^+^). In the latter condition, the activity measured in the presence of **4b** and ADAMTS-5 alone was taken as 100%.

### Statistics

Data are presented as mean ± SEM of at least three independent experiments and were analyzed by GraphPad Prism Software. Statistical analysis was performed using Mann-Whitney test. p<0.05 was considered significant.

### In silico studies

#### Molecular Modelling

The crystal structure of human ADAMTS-5 Mp/Dis domains (PDB code 2RJQ)^**13**^ complexed with its reference inhibitor was minimized using AMBER16 software and ff14SB force field at 300 K, after removing all hydrogen atoms. The complex was placed in a rectangular parallelepiped water box, an explicit solvent model for water, TIP3P, was used and the complex was solvated with a 10 Å water cap. Sodium ions were added as counter-ions to neutralize the system. Two steps of minimization were then carried out; in the first stage, we kept the protein fixed with a position restraint of 500 kcal/mol Å^2^ and we solely minimized the positions of the water molecules. In the second stage, we minimized the entire system through 5000 steps of steepest descent followed by conjugate gradient (CG) until a convergence of 0.05 kcal/Å•mol. Molecular docking calculations were performed with AUTODOCK 4.2 using the improved force field.^**43,44**^ Autodock Tools were used to identify the torsion angles in the ligand, add the solvent model and assign the Kollman atomic charges to the protein, while ligand charges were calculated with the Gasteiger method. A grid spacing of 0.375 Å and a distance-dependent function of the dielectric constant were used for the energetic map calculations. Compound **4b** was subjected to a robust docking procedure already used in virtual screening and pose prediction studies.^**45–47**^ The docked compound was subjected to 200 runs of the AUTODOCK search using the Lamarckian Genetic Algorithm performing 10 000 000 steps of energy evaluation. The number of individuals in the initial population was set to 500 and a maximum of 10 000 000 generations were simulated during each docking run. All other settings were left as their defaults and the best docked conformation was considered. For the modelling of 4b/GM6001/ADAMTS-5 co-complex, GM6001 was first subjected to the docking procedure as above. The so-obtained ADAMTS-5-GM6001 complex was used for the docking evaluation of compound **4b** by using all parameters described above. The results were then clustered by applying a root-mean-square deviation (RMSD) of 6.0 Å. The clusters with a population of at least 40 poses (corresponding to the 20% of the total poses) were considered. The cluster analysis suggested three possible binding orientations for **4b** (**Figure 2-Figure supplement 1**), which were subjected to MD simulations.

#### MD simulations

All simulations were performed using AMBER, version 16. MD simulations were carried out using the ff14SB force field at 300 K. The complex was placed in a rectangular parallelepiped water box. An explicit solvent model for water, TIP3P, was used, and the complex was solvated with a 20 Å water cap. Sodium ions were added as counter-ions to neutralize the system. Prior to MD simulations, two steps of minimization were carried out using the same procedure described above. Particle mesh Ewald (PME) electrostatics and periodic boundary conditions were used in the simulation. The MD trajectory was run using the minimized structure as the starting conformation. The time step of the simulations was 2.0 fs with a cut-off of 10 Å for the non-bonded interactions, and SHAKE was employed to keep all bonds involving hydrogen atoms rigid. Constant-volume periodic boundary MD was carried out for 3.0 ns, during which the temperature was raised from 0 to 300 K. Then 100 ns of constant pressure periodic boundary MD was carried out at 300 K by using the Monte Carlo barostat with anisotropic pressure scaling for pressure control. All the α carbons of the protein were blocked with a harmonic force constant of 10 kcal/mol•Å^2^. General Amber force field (GAFF) parameters were assigned to the ligand, while partial charges were calculated using the AM1-BCC method as implemented in the Antechamber suite of AMBER 16. A representative docking pose belonging to each of the three cluster of poses (C1-C3) was subjected to a 103 ns MD simulation with explicit water molecules. By analyzing the RMSD of the position of compound **4b** during the simulation with respect to the starting pose, we observed an average RMSD of 6.4 and 7.5 Å for C1 and C3. Considering the high degrees of freedom that characterizes **4b**, these two orientations were considered quite stable. On the other hand, the C2 binding mode was highly unstable (average RMSD: 30.4 Å, **Figure 2- Figure supplement 3**) and therefore discarded.

#### Binding Energy Evaluation

Relative binding free energy evaluations were performed using AMBER 16. The trajectories extracted from the last 100 ns of each simulation were used for the calculation, for a total of 100 snapshots (at time intervals of 1 ns). Van der Waals, electrostatic and internal interactions were calculated with the SANDER module of AMBER 16, whereas the Poisson−Boltzman method was employed to estimate polar energies through the molecular mechanics and Poisson Boltzmann surface area (MM-PBSA) module of AMBER 16 as previously reported.^**48**^ Gas and water phases were represented using dielectric constants of 1 and 80, respectively, while nonpolar energies were calculated with MOLSURF program. The entropic term was considered as approximately constant in the comparison of the ligand-protein energetic interactions. All three binding modes were further analysed through MM-PBSA method. This approach averages the contribution of solvation free energy and gas phase energy for snapshots of the ligand-protein complex and the unbound components extracted from MD trajectories. The results of the MM-PBSA analysis suggested pose C1 as the most favorable binding mode, since it showed an interaction energy (ΔPBSA = −14.7 kcal/mol) that was more than 8 kcal/mol lower than that estimated for the binding mode C2 and C3 (**Figure 2- Table supplement 1**). The results obtained from these analyses suggested the MD-refined C1 pose as the most reliable binding disposition of **4b** within ADAMTS-5.

### Chemical Synthesis

The initially investigated compounds **1** and **2** were prepared as previously reported.^**20**^ For synthesis of compounds **3a,b**, **4a-d** and **5** see supplementary material.

## Additional information

The authors declare no conflict of interest.

## Acknowledgments

This work was supported by the Imperial College European Partners Fund 2018, British Heart Foundation project grant (PG/18/15/33566, PI J.A) and by funding from University of Pisa (PRA_2018_20 “Approcci Target/Multitarget per il disegno e lo sviluppo di Small Molecules per terapie innovative” and “Fondi di Ateneo” 2019 to AR and EN). We thank Dr. Suneel Apte (Cleveland Clinic Lerner Research Institute, Ohio, USA) for kindly providing the V1-5GAG vector.

## SUPPLEMENTARY MATERIAL

### Chemistry

#### Instrumentation

Melting points were determined with a Kofler hot-stage apparatus and are uncorrected. ^1^H NMR spectra were recorded in appropriate solvents with a Bruker Avance III HD 400 spectrometer operating at 400 MHz. ^13^C NMR spectra were recorded with the above spectrometer operating at 100.57 MHz. The assignments were made, when possible, with the aid of DEPT, COSY, HSQC experiments. The first order proton chemical shifts (δ) are referenced to residual solvents and J-values are given in Hz. All reactions were followed by TLC on Kieselgel 60 F254 with detection by UV light and/or with ethanolic 10% phosphomolybdic or sulfuric acid, and heating. Kieselgel 60 (Merck, 230-400 mesh) was used for flash chromatography. Some chromatographic separations were conducted by using the automated system Isolera® Prime (Biotage), equipped with UV detector with variable wavelength (200-400 nm) or using prepacked ISOLUTE Flash Si II cartridges (Biotage). Microwave-assisted reactions were run in an Initiator+ (Biotage) microwave synthesizer. All reactions involving air- or moisture-sensitive reagents were performed under an argon atmosphere using anhydrous solvents. Anhydrous dimethylformamide (DMF), dichloromethane (CH_2_Cl_2_), 1,2-dichloethane (DCE) and THF were purchased from Sigma-Aldrich. MgSO_4_ or Na_2_SO_4_ were used as the drying agents for solutions. Elemental analysis has been used to determine the purity of target compounds. Analytical results are within ± 0.40% of the theoretical values.

##### General strategy

The initially investigated compounds **1** and **2** (**Table 1**) were prepared as previously reported.^**1**^ The new compounds **3a,b**, **4a-d** and **5** tested in this study were synthesized as described in **Schemes 1-2**. For this, the alkynyl precursors **10a-d** were prepared as shown in **Scheme 1**. Sulfonyl chlorides **6 b-d**^*1,2*^ were respectively converted into sulfonamides **7b-d**, by reaction with (+/−)-*sec*-butylamine in a mixture H_2_O-dioxane (1:1 *v/v*) and triethyamine (TEA). The known carboxylic acid **8**^**1**^ was protected by treatment with benzyl bromide and caesium carbonate to give benzyl ester **9.** Sulfonamides **7b-d** and **9** were *N*-alkylated by SN2 reaction with propargyl bromide in DMF using potassium carbonate as base to afford alkynes **10a-d** in 57-100 % yields.

**Scheme 1.**
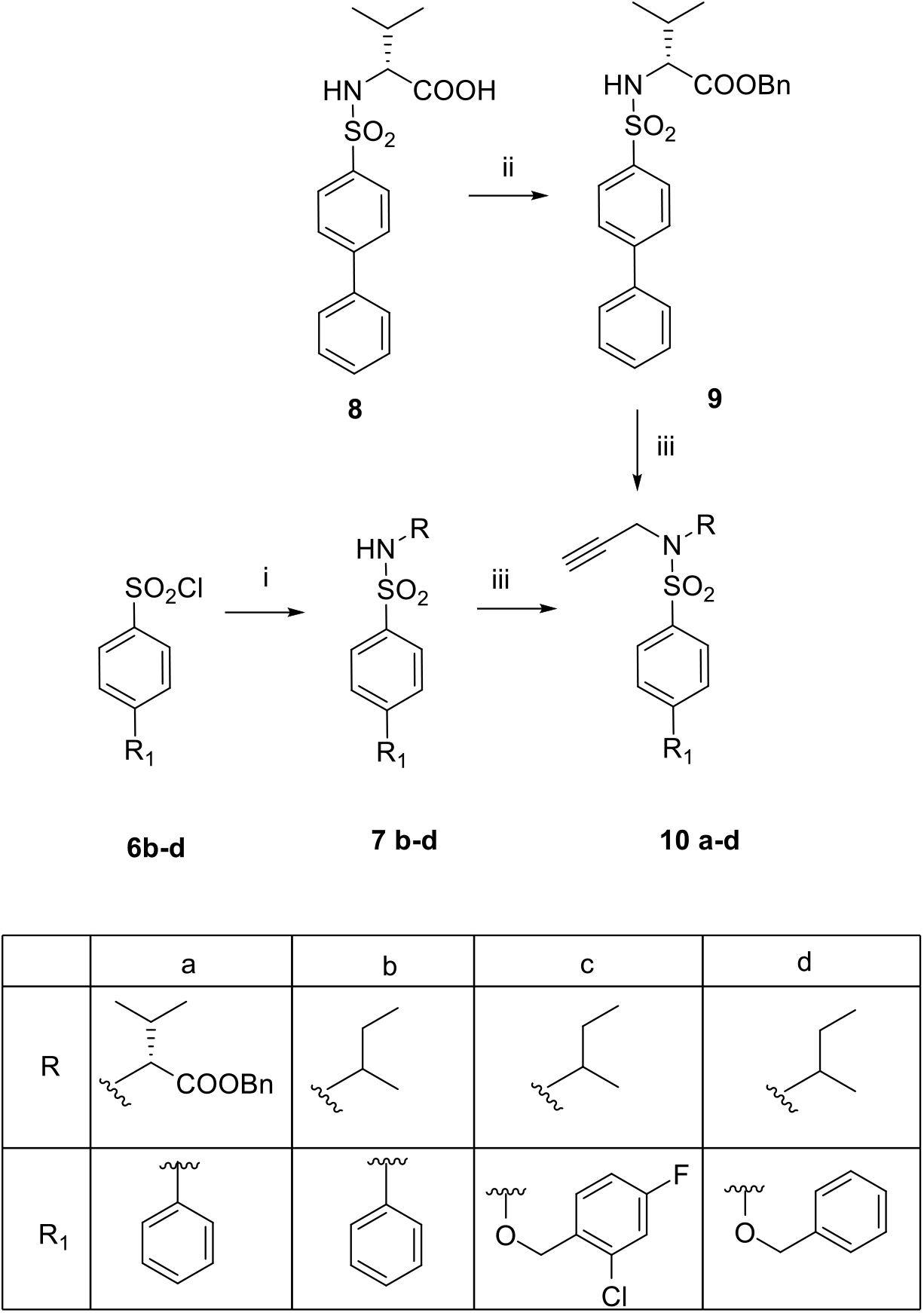
Synthesis of alkynyl precursors **10a-d**. Reagents and conditions: i) (+/−)-*sec*-butylamine, Et_3_N, 1:1 H_2_O-Dioxane, 18 h (**7b**: quantitative yield; **7c**: quantitative yield; **7d**: 30%); ii) BnBr, Cs_2_CO_3_, DMF, 18 h (63%); iii) propargyl bromide, K_2_CO_3_, DMF, 48 h (**10a**: 94%; **10b**: quantitative yield; **10c**: 87%; **10d**: 57%).

The known 3-azidopropyl benzoate **13**^**3**^ was prepared starting from 3-hydroxypropyl *p*-toluenesulfonate^**4**^ (**Scheme 2**). The benzoylation of the hydroxyl portion was achieved by treatment with benzoyl chloride in DCM using DMAP and triethylamine as bases, to give tosylate **16**. 3-Azidopropyl benzoate **13** was obtained by conversion of **16** into the corresponding azide through an S_N_2 reaction (NaN_3_, DMF) in a nearly quantitative yield (98 %).

The β-glycosyl azides **11, 12** and **13** were conjugated to the proper alkynes **10a-d** (**Scheme 2**) by CuAAC click chemistry according to reported conditions.^**1**^ The reactions were performed in a mixture DMF-H_2_O (4:1 *v/v*) with copper(II) sulfate, sodium ascorbate catalytic system and heated under microwave irradiation at 80 °C for 30-45 min. Compound **5** was isolated by flash chromatography in good yield (79%). The desired 1,2,3-triazole derivatives **14a**, **14b** and **15a-d**, were used directly without further purification in the following reaction step.

**Scheme 2.**
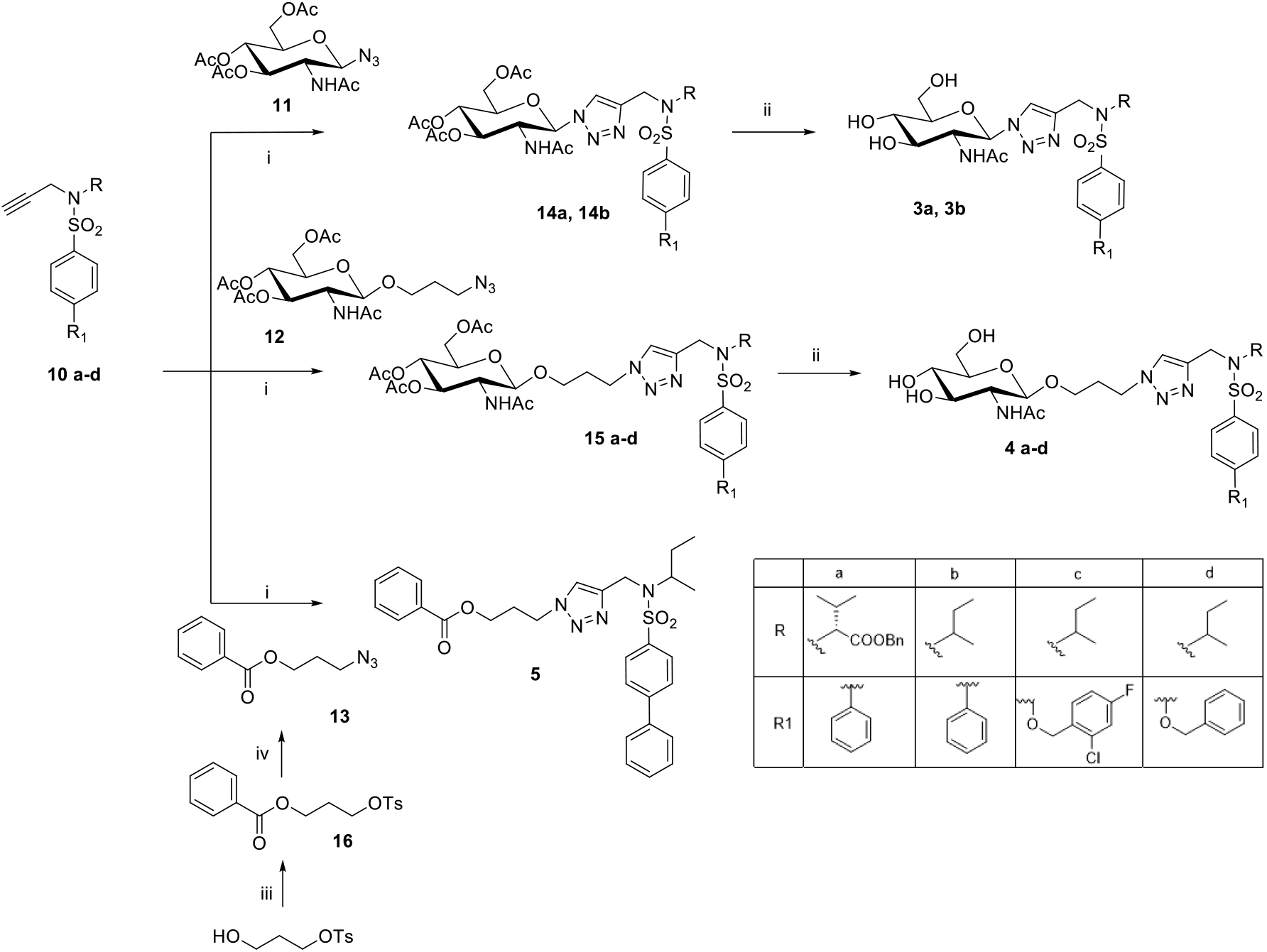
Synthesis of 1,2,3-triazole glycoconjugates **3a**, **3b, 4a-d** and benzoyl derivative **5**. Reagents and conditions for **3a**, **3b, 4a-d**: i) CuSO_4_·5H_2_O, sodium ascorbate, 4:1 DMF-H_2_O, microwave, 80 °C, 30-45 min; ii) NH_3_-MeOH 3.5N, 20-24 h. Yield over two steps: 29% for **3a**; 55% for **3b**; 50% for **4a**; 40% for **4b**; 14% for **4c**, 91% for **4d.** Reagents and conditions for **5:** i) CuSO_4_·5H_2_O, sodium ascorbate, 4:1 DMF-H_2_O, microwave, 80 °C, 45 min (79%); iii) BzCl, Et_3_N, DMAP, DCM room temperature (RT) overnight (o/n) (56%); iv) NaN_3,_ DMF, 80°C, 3h (98%).

Finally, the *O*-deacetylation of **14a, 14b** and **15a-d** by treatment with NH_3_-MeOH 3.5N afforded the deprotected derivatives **3a**, **3b** and **4a-d** in good yields (14-91 %). All compound structures were confirmed by mono- and two-dimensional NMR analyses (^1^H, ^13^C, COSY, HSQC).

#### Synthesis of compounds 11 and 12

Compounds **11** and **12** were prepared as previously reported.^**1**^

#### General Procedure for the Synthesis of Sulfonamides 7 b-d

To a solution of (+/−)-*sec*-butylamine (1 equiv.) in H_2_O (0.8 mL/mmol) and dioxane (0.8 mL/mmol) containing TEA (2 equiv.), the proper sulfonyl chloride (1.2 equiv.) was added. The mixture was stirred at room temperature overnight. Dioxane was evaporated at diminished pressure, and the residue was taken up in H_2_O and extracted with EtOAc (3×10 mL). Organic layers were collected, dried over Na_2_SO_4_, filtered and concentrated in *vacuo*, affording the desired sulfonamides as yellowish solids.

##### N-[(+/−)-sec-butyl]-[1,1’-biphenyl]-4-sulfonamide (7b)

The title compound was prepared from commercial biphenyl-4-sulfonyl chloride **6b** following the general procedure. Yellow solid (quantitative yield). ^1^H NMR (400 MHz, CDCl_3_) δ: 7.94 (m, 2H, Ar-*H*), 7.72 (m, 2H, Ar-*H*), 7.69 (m, 2H, Ar-*H*), 7.50-7.41 (m, 3H, Ar-*H*), 4.49 (bs, 1H, NH), 3.31 ( m, 1H, C*H*CH_3_), 1.43 (m, 2H, C*H*_*2*_CH_3_), 1.06 (d, 3H, *J*_*vic*_=6.8 Hz, CHC*H*_*3*_), 0.82 (t, 3H, *J*_*vic*_=7.2 Hz, CH_2_C*H*_*3*_).

##### N-[(+/−)-sec-butyl]-4-[(2-chloro-4-fluorobenzyl)-oxy]-benzenesulfonamide (7c)

The title compound was prepared from the known sulfonyl chloride **6c**^**2**^ following the general procedure. Yellow solid (quantitative yield). ^1^H NMR (400 MHz, CDCl_3_) δ: 7.83 (m, 2H, Ar-*H*), 7.50 (dd, 1H, *J*=6.0 Hz, *J*=8.6 Hz, Ar-*H*), 7.19 (dd, 1H, *J*=8.4 Hz, *J*=2.6 Hz, Ar-*H*), 7.07-6.96 (m, 3H, Ar-*H*), 5.17 (s, 2H, C*H*_*2*_O), 4.19 (d, 1H, *J*=8.0 Hz, NH), 3.28-3.23 (m, 1H, C*H*CH_3_), 1.44-1.36 (m, 2H, C*H*_*2*_CH_3_), 1.04 (d, 3H, *J*_*vic*_=6.4 Hz, CHC*H*_*3*_), 0.80 (t, 3H, *J*_*vic*_=7.2 Hz, CH_2_C*H*_*3*_).

##### 4-(benzyloxy)-N-[(+/−)-sec-butyl]benzenesulfonamide (7d)

The title compound was prepared from the known sulfonyl chloride **6d**^**2**^ following the general procedure. The crude was purified by flash chromatography (5:1 *n*-hexane-EtOAc) using an Isolute Flash Si II cartridge to afford **7d** as a yellow solid (30% yield). ^1^H NMR (400 MHz, CDCl_3_) δ: 7.80 (m, 2H, Ar-*H*), 7.43-7.40 (m, 5H, Ar-*H*), 7.04 (m, 2H, Ar-*H*), 5.13 (s, 2H, C*H*_*2*_O), 4.13 (d, 1H, *J*=8.4 Hz, NH), 3.25-3.22 (m, 1H, C*H*CH_3_), 1.42-1.38 (m, 2H, C*H*_*2*_CH_3_), 1.03 (d, 3H, *J*_*vic*_=6.4 Hz, CHC*H*_*3*_), 0.80 (t, 3H, *J*_*vic*_=7.2 Hz, CH_2_C*H*_*3*_).

##### (R)-benzyl 2-([1,1’-biphenyl]-4-ylsulfonamido)-3-methylbutanoate (9)

To a solution of the known carboxylic acid **8**^**1**^ (965 mg, 2.895 mmol) in dry DMF (3.8 mL), Cs_2_CO_3_ (707 mg, 2.171 mmol) was added at 0 °C under inert atmosphere (Ar). The mixture was stirred for 1 h at 0 °C and then benzyl bromide (0.26 mL, 2.171 mmol) was added. After stirring overnight at room temperature, the reaction mixture was taken up with water and extracted with EtOAc (3×125 mL). The collected organic layers were washed with brine, dried over Na_2_SO_4_, and concentrated at diminished pressure. The crude was purified by flash chromatography (8:1 *n*-hexane-EtOAc) affording the benzyl derivative **9** as a white solid (577 mg, 63% yield). ^1^H NMR (400 MHz, CDCl_3_) δ: 7.87 (m, 2H, Ar-*H*); 7.64 (m, 2H, Ar-*H*); 7.59-7.57 (m, 2H, Ar-*H*); 7.51-7.41 (m, 3H, Ar-*H*), 7.27-7.24 (m, 3H, Ar-*H*); 7.15-7.13 (m, 2H, Ar-*H*), 5.16 (d, 1H, *J*= 10.1 Hz, N*H*), 4.88, 4.83 (AB system, 2H, *J*_A,B_= 10.8 Hz, CH_2_Ph), 3.84 (dd, 1H, *J*= 4.9 Hz, C*H*N), 2.11-2.06 (m, 1H, CHC*H*_*3*_), 0.98 (d, 3H, *J*_*vic*_=6.8 Hz, C*H*_*3*_), 0.86 (d, 3H, *J*_*vic*_=6.8 Hz, C*H*_*3*_).

#### General procedure for the synthesis of propargyl derivatives 10a-d

To a solution of the proper sulfonamide (1 equiv.) in dry DMF (1.6 mL/mmol) propargyl bromide (80% in toluene, 1.2 equiv.) and potassium carbonate (10 equiv.) were added. The resulting suspension was stirred at room temperature for 2 days under argon atmosphere. The mixture was diluted with water and extracted with EtOAc (3×30 mL). The combined organic phases were washed with brine, dried over Na_2_SO_4_ and the solvent was removed at diminished pressure. The crude product was purified by column chromatography on silica gel or by trituration to give the desired compounds **10a-d**.

##### (R)-Benzyl 3-methyl-2-(N-(prop-2-yn-1-yl)-[1,1’-biphenyl]-4-ylsulfonamido)butanoate (10a)

The title compound was prepared from sulfonamide **9** following the general procedure. The crude product was purified by trituration with *n*-hexane to afford **10a** as a yellow solid (94% yield). ^1^H NMR (400 MHz, CDCl_3_) δ: 7.94-7.88 (m, 2H, Ar-*H*), 7.60-7.55 (m, 4H, Ar-*H*), 7.50-7.40 (m, 3H, Ar-*H*), 7.29-7.26 (m, 3H, Ar-*H*), 7.22-7.19 (m, 2H, Ar-*H*), 5.00, 4.84 (AB system, 2H *J*_A,B_=12.4 Hz, CH_2_Ph), 4.40 (dd, 1H, *J*_gem_=21.2 Hz, *J*= 2.4 Hz, CH_2_C≡), 4.18 (dd, 1H, CH_2_C≡), 4.21-4.05 (m, 1H, C*H*N), 2.26-2.22 (m, 1H, C*H*Me_2_), 2.14 (t, 1H, *J*= 2.4 Hz, C≡C*H*), 1.07 (d, 3H, *J*_*vic*_=6.8 Hz, C*H*_*3*_), 0.95 (d, 3H, *J*_*vic*_=6.8 Hz, C*H*_*3*_).

##### N-[(+/−)-sec-butyl]-N-(prop-2-yn-1-yl)-[1,1’-biphenyl]-4-sulfonamide (10b)

The title compound was prepared from sulfonamide **7b** following the general procedure. The crude product was purified by trituration with *n*-hexane to afford **10b** as a yellow solid (quantitative yield). ^1^H NMR (400 MHz, CDCl_3_) δ: 7.96 (m, 2H, Ar-*H*), 7.70 (m, 2H, Ar-*H*), 7.62 (m, 2H, Ar-*H*), 7.48 (m, 2H, Ar-*H*), 7.45-7.40 (m, 1H, Ar-*H*), 4.10 (dd, 1H, *J*_gem_=18.4 Hz, *J*= 2.4 Hz, CH_2_C≡), 4.00 (dd, 1H, CH_2_C≡), 3.92-3.87 ( m, 1H, C*H*CH_3_), 2.15 (t, 1H, *J*= 2.4 Hz, C≡C*H*), 1.61-1.47 (m, 2H, C*H*_*2*_CH_3_), 1.11 (d, 3H, *J*_*vic*_=6.8 Hz, CHC*H*_*3*_), 0.85 (t, 3H, *J*_*vic*_=7.2 Hz, CH_2_C*H*_*3*_).

##### N-[(+/−)-sec-butyl]-4-[(2-chloro-4-fluorobenzyl)oxy]-N-(prop-2-yn-1-yl)benzenesulfonamide (10c)

The title compound was prepared from sulfonamide **7c** following the general procedure. The crude product was purified by trituration with *n*-hexane to afford **10c** as a yellow solid (87% yield). ^1^H NMR (400 MHz, CDCl_3_) δ: 7.86 (m, 2H, Ar-*H*), 7.50 (dd, 1H, *J*=5.9 Hz, *J*=8.4 Hz, Ar-*H*), 7.18 (dd, 1H, *J*=8.4 Hz, *J*=2.5 Hz, Ar-*H*), 7.06-7.00 (m, 3H, Ar-*H*), 5.17 (s, 2H, C*H*_*2*_O), 4.10 (dd, 1H, *J*_gem_=18.5 Hz, *J*= 2.4 Hz, CH_2_C≡), 4.05 (dd, 1H, CH_2_C≡), 3.86-3.80 (m, 1H, C*H*CH_3_), 2.15 (t, 1H, *J*= 2.4 Hz, C≡C*H*), 1.61-1.45 (m, 2H, C*H*_*2*_CH_3_), 1.09 (d, 3H, *J*_*vic*_=6.4 Hz, CHC*H*_*3*_), 0.83 (t, 3H, *J*_*vic*_=7.2 Hz, CH_2_C*H*_*3*_).

##### 4-(Benzyloxy)- N-[(+/−)-sec-butyl]-N-(prop-2-yn-1-yl)benzenesulfonamide (10d)

The title compound was prepared from sulfonamide **7d** following the general procedure. The crude product was purified by flash chromatography (10:1 *n*-hexane-EtOAc) using an Isolute Flash Si II cartridge to afford **10c** as a yellow solid (57% yield). ^1^H NMR (400 MHz, CDCl_3_) δ: 7.83 (m, 2H, Ar-*H*), 7.43-7.34 (m, 5H, Ar-*H*), 7.02 (m, 2H, Ar-*H*), 5.12 (s, 2H, C*H*_*2*_O), 4.09 (dd, 1H, *J*_gem_=18.4 Hz, *J*= 2.4 Hz, CH_2_C≡), 4.04 (dd, 1H, CH_2_C≡), 3.85-3.80 (m, 1H, C*H*CH_3_), 2.11 (t, 1H, *J*= 2.4 Hz, C≡C*H*), 1.57-1.45 (m, 2H, C*H*_*2*_CH_3_), 1.09 (d, 3H, *J*_*vic*_=6.8 Hz, CHC*H*_*3*_), 0.83 (t, 3H, *J*_*vic*_=7.2 Hz, CH_2_C*H*_*3*_).

#### General procedure for the synthesis of glycoconjugates 3a, 3b, 4a-d and benzoyl derivative 5

The appropriate alkyne (1 equiv.) and the opportune azide (1.1 equiv.) were dissolved in a mixture of 4:1 DMF-H_2_O (24 mL/ mmol) in the presence of CuSO_4_·5H_2_O (1.5 equiv.) and sodium ascorbate (3 equiv.). The solution was heated under microwave irradiation to 80 °C for 30-45 min, then diluted with Et_2_O (12 mL) and washed with saturated aq NH_4_Cl (25 mL). The organic phase was separated and the aq layer extracted with Et_2_O (2×25 mL). The collected organic extracts were dried (Na_2_SO_4_), filtered and concentrated under diminished pressure. Flash chromatographic purification over silica gel of the crude product gave triazole-linked derivative **5** pure as a white solid. The acetylated glycoconjugates **14a**, **14b** and **15a-d** were used in the following step without any purification. A solution of the appropriate acetylated glycoconjugate (**14a**, **14b** or **15a-d**, mmol) in MeOH (1.0 mL) was treated with NH_3_-MeOH 7N (1.0 mL) and the solution was stirred at room temperature until the starting compound was completely reacted (TLC, 8:2 CHCl_3_-MeOH, 20-24 h). The solution was co-evaporated with toluene (4×10 mL) under diminished pressure. Flash chromatographic purification over silica gel of the crude product gave pure triazole-linked derivatives (**3a**, **3b** or **4a-d**) as white solids.

##### Glycoconjugate 3a

The title compound was obtained from alkyne **10a** and azide **11** following the general procedure. After treatment, the glycoconjugate **14a** was isolated and used directly in the basic hydrolysis with NH_3_-MeOH 3.5N. The crude was purified by flash chromatography (20:1 CHCl_3_-MeOH) using an Isolute Flash Si II cartridge to afford **3a** as a white solid (29% yield from **10a**); mp 114-116 °C; ^1^H NMR (400 MHz, CD_3_OD) δ: 8.18 (s, 1H, Ar-*H* triazole), 7.92-7.80 (m, 2H, Ar-*H*), 7.70-7.63 (m, 4H, Ar-*H*), 7.51-7.37 (m, 3H, Ar-*H*), 7.25-7.18 (m, 5H, Ar-*H*), 5.79 (d, 1H, *J*_1,2_=9.8 Hz, H-1), 4.89, 4.72 (AB system, 2H, *J*_A,B_=12.2 Hz, PhCH_2_O), 4.84, 4.66 (AB system, 2H, *J*_A,B_=16.6 Hz, CH_2_NSO_2_), 4.23 (bt, 1H, *J*_1,2_=*J*_2,3_=9.8 Hz, H-2), 4.09 (d, 1H, *J*_vic_=10.6 Hz, C*H*N), 3.91 (bd, 1H, *J*_6a,6b_=12.1 Hz, H-6b), 3.78-3.67 (m, 2H, H-3, H-5), 3.63-3.51 (m, 2H, H-4, H-6a), 2.29 (m, 1H, C*H*Me_2_), 1.79 (s, 3H, C*H*_*3*_CON), 0.85 (d, 3H, *J*_*vic*_=6.5 Hz, C*H*_*3*_), 0.71 (d, 3H, *J*_*vic*_=6.5 Hz, C*H*_*3*_); ^13^C NMR (100 MHz, CD_3_OD) δ: 173.4, 171.7 (2×C=O), 147.0 (Ar-*C*-SO_2_, *C*-triazole), 140.3, 139.2, 136.5 (3×Ar-*C*), 1130.2-125.3 (Ar-*C*H), 125.3 (CH-triazole), 88.2 (C-1), 81.4 (C-5), 75.8 (C-3), 71.4 (C-4), 67.7 (O*C*H_2_Ph), 67.5 (CHN), 62.4 (C-6), 56.9 (C-2), 40.9 (CH_2_NSO_2_), 30.2 (*C*HMe_2_), 22.6 (MeCON), 20.3, 19.5 (*Me*_2_CH). Elemental analysis calcd (%) for C_35_H_41_N_5_O_9_S: C 59.39, H 5.84, N 9.89; found: C 59.42, H 5.90, N 9.93.

##### Glycoconjugate 3b

The title compound was obtained from alkyne **10b** and azide **11** following the general procedure. After treatment, the glycoconjugate **14b** was isolated and used directly in the basic hydrolysis with NH_3_-MeOH 3.5N. The crude was purified by flash chromatography (20:1 CHCl_3_-MeOH) using an Isolute Flash Si II cartridge to afford **3b** as a white solid (55% yield from **10b**); mp 211-213 °C. The NMR analysis of **3b** (CD_3_OD) showed a mixture of the two diastereoisomeric forms in the ratio of 1:1, measured on the relative intensities of the H-1 signals at δ 5.79 and 5.78 respectively. ^1^H NMR (400 MHz, CD_3_OD) *one diastereoisomeric form* δ: 8.08 (s, 1H, Ar-*H* triazole), 5.79 (d, 1H, *J*_1,2_=9.8 Hz, H-1), 4.52, 4.40 (AB system, 2H, *J*_A,B_= 16.6 Hz, C*H*_*2*_NSO_2_), 1.78 (s, 3H, C*H*_*3*_CONH), 1.57-1.42 (m, 2H, CH_3_C*H*_*2*_), 0.99 (d, 3H, *J*_*vic*_=6.7 Hz, C*H*_*3*_CH), 0.72 (t, 3H, *J*_*vic*_=7.4 Hz, C*H*_*3*_CH_2_); *other diastereoisomeric form* δ: 8.07 (s, 1H, Ar-*H* triazole), 5.78 (d, 1H, *J*_1,2_=9.8 Hz, H-1), 4.48, 4.43 (AB system, 2H, *J*_A,B_=17.1 Hz, C*H*_*2*_NSO_2_), 1.77 (s, 3H, C*H*_*3*_CONH), 1.41-1.31 (m, 2H, CH_3_C*H*_*2*_), 0.98 (d, 3H, *J*_*vic*_=6.7 Hz, C*H*_*3*_CH), 0.70 (t, 3H, *J*_*vic*_=7.4 Hz, C*H*_*3*_CH_2_); *cluster of signals for both diastereoisomeric forms* δ: 7.93-7.90 (m, 2H, Ar-*H*), 7.84-7.82 (m, 2H, Ar-*H*), 7.71-7.69 (m, 2H, Ar-*H*), 7.51-7.47 (m, 2H, Ar-*H*), 7.43-7.39 (m, 1H, Ar-*H*), 4.25-4.19 (m, 1H, C*H*N), 3.92-3.84 (m, 2H, H-2, H-6b), 3.76-3.71 (m, 1H, H-6a), 3.68 (bt, 1H, *J*_*2,3*_= *J*_*3,4*_=9.3 Hz, H-3); 3.60-3.51 (m, 2H, H-4, H-5). ^13^C NMR (100 MHz, CD_3_OD) *one diastereoisomeric form* δ: 124.8 (*C*H-triazole), 88.4 (C-1), 76.2 (C-3), 57.8 (*C*HN), 57.0 (C-2), 38.9 (*C*H_2_NSO_2_), 29.4 (CH_3_*C*H_2_), 22.9 (*C*H_3_CON), 19.1 (*C*H_3_CH), 11.9 (*C*H_3_CH_2_); *other diastereoisomeric form* δ: 124.6 (*C*H-triazole), 88.3 (C-1), 76.1 (C-3), 57.7 (*C*HN), 56.9 (C-2), 38.8 (*C*H_2_NSO_2_), 29.3 (CH_3_*C*H_2_), 22.8 (*C*H_3_CON), 18.9 (*C*H_3_CH), 11.8 (*C*H_3_CH_2_); *cluster of signals for both diastereoisomeric forms* δ: 173.5 (C=O), 147.3-147.0 (Ar-*C*-SO_2_, *C*-triazole), 141.1-140.8 (2×Ar-*C*), 130.4-128.5 (Ar-*C*H), 81.6 (C-5), 71.6 (C-4), 62.6 (C-6). Elemental analysis calcd (%) for C_27_H_35_N_5_O_7_S: C 56.53, H 6.15, N 12.21; found: C 56.55, H 6.17, N 12.23.

##### Glycoconjugate 4a

The title compound was obtained from alkyne **10a** and azide **12** following the general procedure. After treatment, the glycoconjugate **15a** was isolated and used directly in the basic hydrolysis with NH_3_-MeOH 3.5N. The crude was purified by flash chromatography (20:1 CHCl_3_-MeOH) using an Isolute Flash Si II cartridge to afford **4a** as a white solid (50% yield from **10a**); mp 88-90 °C; ^1^H NMR (400 MHz, CD_3_OD) δ: 7.89 (s, 1H, Ar-*H* triazole), 7.84-7.81 (m, 2H, Ar-*H*), 7.67-7.63 (m, 4H, Ar-*H*), 7.52-7.43 (m, 3H, Ar-*H*), 7.27-7.00 (m, 5H, Ar-*H*), 4.94, 4.81(AB system, 2H, *J*_A,B_= 12.2 Hz, C*H*_*2*_Ph), 4.82, 4.66 (AB system, 2H, *J*_A,B_=16.6 Hz, C*H*_*2*_NSO_2_), 4.50-4.40 (m, 2H, C*H*_*2*_N), 4.36 (d, 1H, *J*_1,2_=8.4 Hz, H-1), 4.16 (d, 1H, *J*_vic_=10.6 Hz, C*H*N), 3.92-3.82 (m, 2H, H-6b, ½C*H*_*2*_O), 3.73-3.66 (m, 2H, H-2, H-6a), 3.48-3.33 (m, 3H, H-3, H-4, ½C*H*_*2*_O), 3.26 (ddd, 1H, *J*_5,6b_=2.6 Hz, *J*_5,6a_=6.0 Hz, *J*_4,5_=10.0 Hz, H-5), 2.08-2.02 (m, 2H, CH_2_), 2.27-2.25 (m, 1H, C*H*CH_3_), 2.01 (s, 3H, C*H*_*3*_CONH), 0.87 (d, 3H, *J*=6.5 Hz, C*H*_*3*_), 0.77 (d, 3H, *J*=6.5 Hz, C*H*_*3*_); ^13^C NMR (100 MHz, CD_3_OD) δ:173.9, 171.9 (2×C=O), 147.1 (Ar-*C*-SO_2_, *C*-triazole), 140.4, 139.7, 136.8 (3×Ar-*C*), 130.4-128.5 (Ar-*C*H), 126.5 (*C*H-triazole), 103.1 (C-1), 78.2 (C-5), 76.2 (C-3), 72.3 (C-4), 68.0 (PhCH_2_O), 67.5 (CHN), 66.8 (CH_2_O), 63.0 (C-6), 57.6 (C-2), 48.4 (CH_2_N), 41.3 (CH_2_NSO_2_), 31.8 (CH_2_), 30.2 (Me_2_*C*H), 23.4 (*C*H_3_CON), 20.4, 19.8 (*Me*_2_CH). Elemental analysis calcd (%) for C_38_H_47_N_5_O_10_S: C 59.59, H 6.19, N 9.14; found: C 59.61, H 6.17, N 9.20.

##### Glycoconjugate 4b

The title compound was obtained from alkyne **10b** and azide **12** following the general procedure. After treatment, the glycoconjugate **15b** was isolated and used directly in the basic hydrolysis with NH_3_-MeOH 3.5N. The crude was purified by flash chromatography (20:1 CHCl_3_-MeOH) using an Isolute Flash Si II cartridge to afford **4b** as a white solid (40% yield from **10b**); mp 73-75 °C. The NMR analysis of **4b** (CD_3_OD) showed a mixture of the two diastereoisomeric forms in the ratio of 1:1, measured on the relative intensities of the C*H*_*3*_CH signals at δ 1.01 and 1.00 respectively. ^1^H NMR (400 MHz, CD_3_OD) *one diastereoisomeric form* δ: 7.98 (s, 1H, Ar-*H* triazole), 2.03 (s, 3H, C*H*_*3*_CONH), 1.55-1.45 (m, 2H, CH_3_C*H*_*2*_), 1.01 (d, 3H, *J*_*vic*_=6.7 Hz, C*H*_*3*_CH), 0.71 (t, 3H, *J*_*vic*_=7.3 Hz, C*H*_*3*_CH_2_); *other diastereoisomeric form* δ: 7.97 (s, 1H, Ar-*H* triazole), 2.02 (s, 3H, C*H*_*3*_CONH), 1.44-1.35 (m, 2H, CH_3_C*H*_*2*_), 1.00 (d, 3H, *J*_*vic*_=6.7 Hz, C*H*_*3*_CH), 0.70 (t, 3H, *J*_*vic*_=7.3 Hz, C*H*_*3*_CH_2_); *cluster of signals for both diastereoisomeric forms* δ: 7.93-7.90 (m, 2H, Ar-*H*), 7.84-7.82 (m, 2H, Ar-*H*), 7.71-7.69 (m, 2H, Ar-*H*), 7.51-7.48 (m, 2H, Ar-*H*), 7.44-7.40 (m, 1H, Ar-*H*), 4.56-4.38 (m, 4H, C*H*_*2*_N, C*H*_*2*_NSO_2_), 4.36 (d, 1H, *J*_1,2_=8.4 Hz, H-1), 3.90-3.80 (m, 3H, H-6b, C*H*N, ½C*H*_*2*_O), 3.73-3.66 (m, 2H, H-2, H-6a), 3.45 (bt, 1H, *J*_2,3_=*J*_3,4_=9.5 Hz, H-3); 3.39-3.27 (m, 2H, H-4, ½C*H*_*2*_O), 3.26 (ddd, 1H, *J*_5,6b_=1.7 Hz, *J*_5,6a_= 5.2 Hz, *J*_4,5_=10.2 Hz, H-5), 2.17-2.08 (m, 2H, C*H*_*2*_). ^13^C NMR (100 MHz, CD_3_OD) *one diastereoisomeric form* δ: 173.9 (C=O), 126.2 (*C*H-triazole), 66.5 (*C*H_2_O), 57.6 (C-2), 31.6 (*C*H_2_), 29.2 (CH_3_*C*H_2_), 18.9 (*C*H_3_CH), 11.7 (*C*H_3_CH_2_); *other diastereoisomeric form* δ: 173.8 (C=O), 126.1 (*C*H-triazole), 66.4 (*C*H_2_O), 57.5 (C-2), 31.5 (*C*H_2_), 29.1 (CH_3_*C*H_2_), 18.7 (*C*H_3_CH), 11.5 (*C*H_3_CH_2_); *cluster of signals for both diastereoisomeric forms* δ: 147.0-146.8 (Ar-*C*-SO_2_, *C*-triazole), 140.9-140.5 (2×Ar-*C*), 130.2-128.3 (Ar-*C*H), 102.8 (C-1), 78.0 (C-5), 76.0 (C-3), 72.1 (C-4), 62.8 (C-6), 57.4 (*C*HN), 48.1 (*C*H_2_N), 38.7 (*C*H_2_NSO_2_), 23.1 (*C*H_3_CON). Elemental analysis calcd (%) for C_30_H_41_N_5_O_8_S: C 57.04, H 6.54, N 11.09; found: C 57.07, H 6.57, N 11.12.

##### Glycoconjugate 4c

The title compound was obtained from alkyne **10c** and azide **12** following the general procedure. After treatment, the glycoconjugate **15c** was isolated and used directly in the basic hydrolysis with NH_3_-MeOH 3.5N. The crude was purif by flash chromatography (CHCl_3_-MeOH 95:1) using Biotage Isolera (5g Zip Sphere Column) to afford **4c** as a white solid (14% yield from **10c**); mp 100-102 °C. The NMR analysis of **4c** (CD_3_OD) showed a mixture of the two diastereoisomeric forms in the ratio of 1:1, measured on the relative intensities of the C*H*_*3*_CH signals at δ 0.97 and 0.96 respectively. ^1^H NMR (400 MHz, CD_3_OD) *one diastereoisomeric form* δ: 7.95 (s, 1H, Ar-*H* triazole), 4.44, 4.38 (AB system, 2H, *J*_A,B_=16.3 Hz, C*H*_*2*_NSO_2_), 2.03 (s, 3H, C*H*_*3*_CONH), 0.97 (d, 3H, *J*_*vic*_=6.7 Hz, C*H*_*3*_CH), 0.67 (t, 3H, *J*_*vic*_=7.3 Hz, C*H*_*3*_CH_2_); *other diastereoisomeric form* δ: 7.94 (s, 1H, Ar-*H* triazole), 4.42, 4.40 (AB system, 2H, *J*_A,B_= 16.5 Hz, C*H*_*2*_NSO_2_), 2.02 (s, 3H, C*H*_*3*_CONH), 0.96 (d, 3H, *J*_*vic*_=6.7 Hz, C*H*_*3*_CH), 0.66 (t, 3H, *J*_*vic*_=7.3 Hz, C*H*_*3*_CH_2_); *cluster of signals for both diastereoisomeric forms* δ: 7.80 (m, 2H, Ar-*H*), 7.60 (dd, 1H, *J*=6.0 Hz, *J*=8.4 Hz, Ar-*H*), 7.30 (dd, 1H, *J*=2.6 Hz, *J*=8.4 Hz, Ar-*H*), 7.17 (m, 2H, Ar-*H*), 7.12 (dd, 1H, *J*=2.6 Hz, *J*=8.4 Hz, Ar-*H*), 5.23 (s, 2H, Ph*CH*_2_O), 4.48 (bt, 2H, *J*_vic_=6.6 Hz, C*H*_*2*_N), 4.38 (d, 1H, *J*_1,2_=8.4 Hz, H-1), 3.93-3.85 (m, 2H, H-6b, ½C*H*_*2*_O), 3.79 (m, 1H, C*H*N), 3.73-3.67 (m, 2H, H-2, H-6a), 3.46 (bt, 1H, *J*_2,3_=*J*_3,4_=9.3 Hz, H-3); 3.40-3.31 (m, 2H, H-4, ½C*H*_*2*_O), 3.26 (ddd, 1H, *J*_5,6b_=2.0 Hz, *J*_5,6a_=5.6 Hz, *J*_4,5_=9.6 Hz, H-5), 2.18-2.08 (m, 2H, C*H*_*2*_), 1.50-1.32 (m, 2H, CH_3_C*H*_*2*_). ^13^C NMR (100 MHz, CD_3_OD) *one diastereoisomeric form* δ: 173.9 (C=O), 126.1 (*C*H-triazole), 66.4 (*C*H_2_O), 57.4 (C-2), 31.5 (*C*H_2_), 29.1 (CH_3_*C*H_2_), 18.8 (*C*H_3_CH), 11.6 (*C*H_3_CH_2_); *other diastereoisomeric form* δ: 173.8 (C=O), 126.0 (*C*H-triazole), 66.3 (*C*H_2_O), 57.3 (C-2), 31.4 (*C*H_2_), 29.0 (CH_3_*C*H_2_), 18.6 (*C*H_3_CH), 11.5 (*C*H_3_CH_2_); *cluster of signals for both diastereoisomeric forms* δ: 163.3 (Ar-*C*O), 147.4-147.3 (Ar-*C*-SO_2_, *C*-triazole), 135.5, 134.5, 131.6 (Ar-*C*), 132.6-130.4 (Ar-*C*H), 118.0-115.2 (Ar-*C*H), 102.8 (C-1), 78.0 (C-5), 76.0 (C-3), 72.1 (C-4), 68.3 (Ph*C*H_2_O), 62.8 (C-6), 57.3 (*C*HN), 23.1 (*C*H_3_CON), 48.1 (*C*H_2_N), 38.6 (*C*H_2_NSO_2_), Elemental analysis calcd (%) for C_31_H_41_ClFN_5_O_9_S: C 52.13, H 5.79, N 9.81; found: C 52.16, H 5.77, N 9.83.

##### Glycoconjugate 4d

The title compound was obtained from alkyne **10d** and azide **12** following the general procedure. After treatment, the glycoconjugate **15d** was isolated and used directly in the basic hydrolysis with NH_3_-MeOH 3.5N. The crude was purified by flash chromatography (20:1 CHCl_3_-MeOH) using an Isolute Flash Si II cartridge to afford **4d** as a white solid (91% yield from **10d**); mp 98-100 °C. The NMR analysis of **4d** (CD_3_OD) showed a mixture of the two diastereoisomeric forms in the ratio of 1:1, measured on the relative intensities of the C*H*_*3*_CH signals at δ 0.96 and 0.95 respectively. ^1^H NMR (400 MHz, CD_3_OD) *one diastereoisomeric form* δ: 7.94 (s, 1H, Ar-*H* triazole), 4.43, 4.37 (AB system, 2H, *J*_A,B_=15.1 Hz, C*H*_*2*_NSO_2_), 2.03 (s, 3H, C*H*_*3*_CONH), 0.96 (d, 3H, *J*_*vic*_=6.7 Hz, C*H*_*3*_CH), 0.67 (t, 3H, *J*_*vic*_=7.3 Hz, C*H*_*3*_CH_2_); *other diastereoisomeric form* δ: 7.93 (s, 1H, Ar-*H* triazole), 4.41, 4.39 (AB system, 2H, *J*_A,B_=15.0 Hz, C*H*_*2*_NSO_2_), 2.02 (s, 3H, C*H*_*3*_CONH), 0.95 (d, 3H, *J*_*vic*_=6.7 Hz, C*H*_*3*_CH), 0.66 (t, 3H, *J*_*vic*_=7.3 Hz, C*H*_*3*_CH_2_); *cluster of signals for both diastereoisomeric forms* δ: 7.79-7.70 (m, 2H, Ar-*H*), 7.45-7.33 (m, 5H, Ar-*H*), 7.19-7.10 (m, 2H, Ar-*H*), 5.18 (s, 2H, Ph*CH*_*2*_O), 4.48 (bt, 2H, *J*_vic_=6.5 Hz, C*H*_*2*_N), 4.45-4.35 (m, 1H, C*H*N), 4.38 (d, 1H, *J*_1,2_=8.5 Hz, H-1), 3.94-3.86 (m, 2H, H-6b, ½C*H*_*2*_O), 3.74-3.61 (m, 2H, H-2, H-6a), 3.46 (bt, 1H, *J*_2,3_=*J*_3,4_=9.5 Hz, H-3); 3.40-3.31 (m, 2H, H-4, ½C*H*_*2*_O), 3.26 (ddd, 1H, *J*_5,6b_=2.0 Hz, *J*_5,6a_=5.7 Hz, *J*_4,5_=9.6 Hz, H-5), 2.15-2.08 (m, 2H, C*H*_*2*_), 1.48-1.33 (m, 2H, CH_3_C*H*_*2*_). ^13^C NMR (100 MHz, CD_3_OD) *one diastereoisomeric form* δ: 126.1 (*C*H-triazole), 66.4 (*C*H_2_O), 62.8 (C-6), 57.4 (C-2), 31.5*C*H(_2_), 29.1 (C_3_H*C*H_2_), 18.8 *C*(H_3_CH), 11.6 (*C*H_3_CH_2_); *other diastereoisomeric form* δ: 126.0 (*C*H-triazole), 66.3 (*C*H_2_O), 62.7 (C-6), 57.3 (C-2), 31.4 (*C*H_2_), 29.0 (CH_3_*C*H_2_), 18.6 (*C*H_3_CH), 11.5 (*C*H_3_CH_2_); *cluster of signals for both diastereoisomeric forms* δ: 173.9 (C=O), 163.6 (Ar-*C*O), 137.8 (Ar-*C*-SO_2_, *C*-triazole), 134.0 (Ar-*C*), 130.3-128.7 (Ar-*C*H), 116.3 (Ar-*C*H), 102.8 (C-1), 78.0 (C-5), 76.0 (C-3), 72.1 (C-4), 71.4 (Ph*C*H_2_O), 23.1 (*C*H_3_CON), 57.2 (*C*HN), 48.1 (*C*H_2_N), 38.6 (*C*H_2_NSO_2_). Elemental analysis calcd (%) for C_31_H_43_N_5_O_9_S: C 56.26, H 6.55, N 10.58; found: C 56.29, H 6.57, N 10.60.

##### Benzoyl derivative 5

The title compound was prepared from alkyne **10b** and azide **13** following the general procedure. After a Flash chromatography using Biotage Isolera (10g Zip Sphere Column, 1:1 EtOAc-*n*-hexane) the benzoyl derivative **5** was isolated as a colourless oil (79% yield). ^1^H NMR (400 MHz, CDCl_3_) δ: 8.08-8.06 (m, 2H, Ar-*H*), 7.91-7.88 (m, 5H, Ar-*H*), 7.87 (s, 1H, Ar-*H* triazole), 7.75-7.73 (m, 2H, Ar-*H*), 7.64-7.60 (m, 5H, Ar-*H*), 7.52-7.43 (m, 5H, Ar-*H*), 4.56 (t, 2H, *J*_vic_=7.1 Hz, C*H*_*2*_N), 4.51-4.41(m, 2H, C*H*_*2*_NSO_2_), 4.37 (t, 2H, *J*_vic_=6.0 Hz, C*H*_*2*_O), 3.92-3.87 (m, 1H, C*H*N), 2.47-2.40 (m, 2H, C*H*_*2*_), 1.55-1.46, 1.45-1.35 (2m, each 1H, CH_3_C*H*_*2*_), 1.02 (d, 3H, *J*_*vic*_=6.8 Hz, C*H*_*3*_CH), 0.71 (t, 3H, *J*_*vic*_=7.3 Hz, C*H*_*3*_CH_2_); ^13^C NMR (100 MHz, CDCl_3_) δ: 166.3 (C=O), 146.6, 145.4 (Ar-*C*-SO_2_, *C*-triazole), 139.5, 139.2, 133.2 (3×Ar-*C*), 129.8-127.3 (Ar-*C*H), 124.1 (*C*H-triazole), 61.4 (*C*H_2_O), 56.3 (*C*HN), 47.3 (*C*H_2_N), 38.2 (*C*H_2_NSO_2_), 29.7 (*C*H_2_), 28.1 (CH_3_*C*H_2_), 18.4 (*C*H_3_CH), 11.1 (*C*H_3_CH_2_). Elemental analysis calcd (%) for C_29_H_32_N_4_O_4_S: C 65.39, H 6.06, N 10.52; found: C 65.42, H 6.07, N 10.56.

#### Synthesis of 3-(tosyloxy)propyl benzoate 16

To a solution of 3-hydroxypropyl *p*-toluenesulfonate^**4**^ (800 mg, 3.47 mmol), DMAP (84 mg, 0.69 mmol) and Et_3_N (0.72 mL, 5.20 mmol) in 8.7 mL of dry DCM, benzoyl chloride (1.39 mL, 12.05 mmol) was added dropwise at 0°C under inert atmosphere (Ar). The reaction mixture was stirred at RT overnight and then quenched with water and extracted with DCM (3×125 mL). The combined organic phases were dried (Na_2_SO_4_), filtered and concentrated under diminished pressure. The crude was purified by flash chromatography (10:1 n-hexane-EtOAc) using an Isolute Flash Si II cartridge to afford **16** as colourless oil (56% yield). ^1^H NMR (400 MHz, CDCl_3_) δ: 7.99-7.88 (m, 2H, Ar-*H*), 7.76-7.71 (m, 2H, Ar-*H*), 7.56-7.52 (m, 1H, Ar-*H*) 7.44-7.38 (m, 2H, Ar-*H*), 7.26-7.23 (m, 2H, Ar-*H*),4.30 (t, 2H, *J*=6.4 Hz, C*H*_*2*_ OCO), 4.19 (t, 2H, , *J*=6.4 Hz*,CH_2_*OSO_2_), 2.15-2.10 (m, 2H, CH_2_).

##### Synthesis of 3-azido propyl benzoate 13^3^

To a solution of **16** (650 mg, 1.94 mmol) in dry DMF (13 mL), NaN_3_ (370 mg, 5.7 mmol) was added. The reaction mixture was refluxed at 80 °C for 3h and cooled to RT. Then, 125 mL of CHCl_3_ were added and the mixture was washed with water (5 × 125 mL). The organic phase was dried (Na_2_SO_4_), filtered and concentrated under diminished pressure, affording compound **13** without any further purification as a yellow solid (390 mg, quantitative yield).^1^H NMR (400 MHz, CDCl_3_) δ: 8.03-8.05 (m, 2H, Ar-*H*), 7.58-7.55 (m, 1H, Ar-*H*), 7.47-7.43 (m, 2H, Ar-*H*),4.43 (t, 2H, *J*=6.4 Hz, C*H*_*2*_ OCO), 3.49 (t, 2H, *J*=6.4 Hz, *CH*_*2*_OSO_2_), 2.06 (m, 2H, *CH*_*2*_).

---

**Figure 1- Figure supplement 1.**
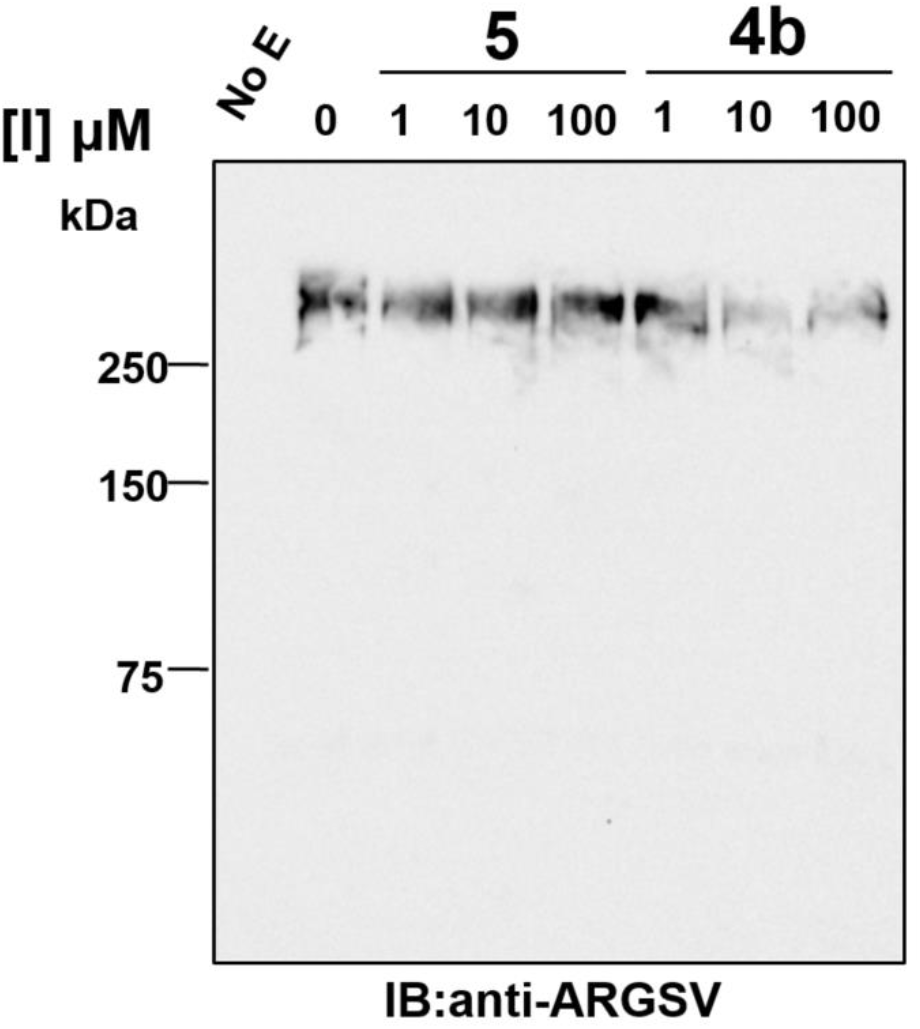
Representative anti-ARGSV western blot. Compounds 4b and 5 were incubated with ADAMTS-5 (1 nM) for 2 h at 37°C before addition of aggrecan (20 μg). Following SDS-PAGE and western blot, fragments cleaved at the Glu392↓Ala393 bond were detected by a monoclonal neoepitope antibody recognizing the new C-terminal fragment (anti-ARGSV) and analyzed by densitometric analysis.

**Figure 2 -Figure supplement 1.**
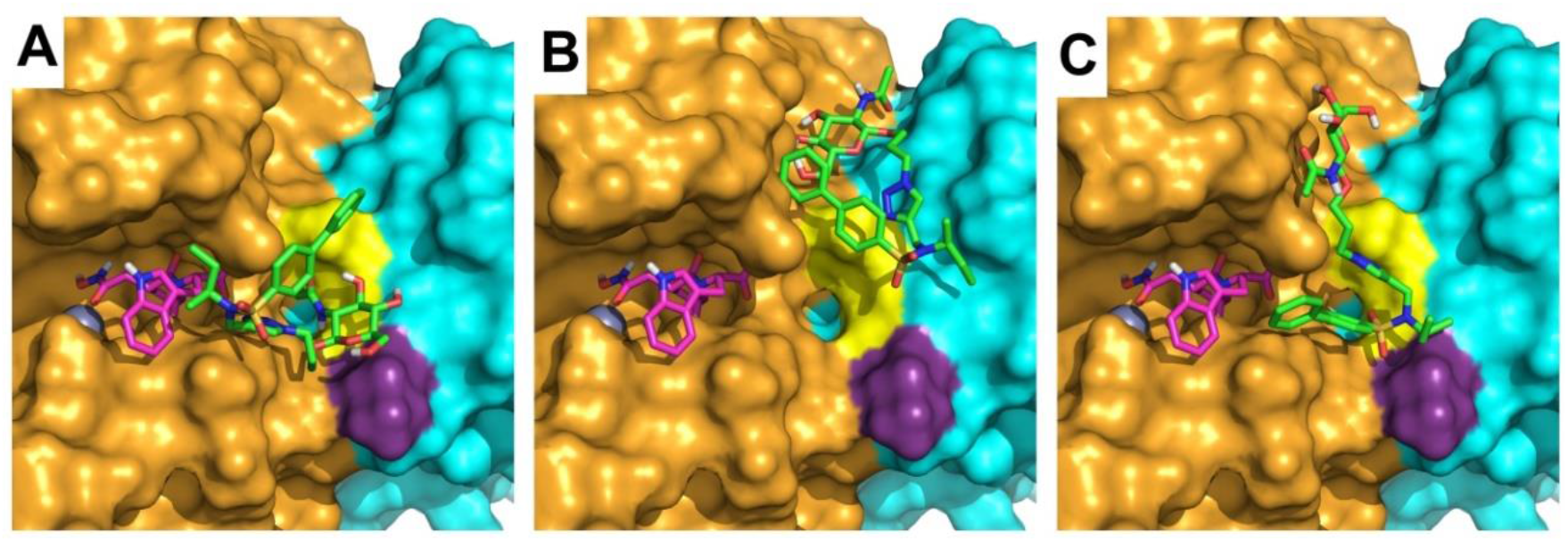
Close-up view of the three binding modes predicted for compound 4b within ADAMTS-5. The representative binding pose of each cluster (poses C1-C3) is shown. The Mp and Dis domains are shown in cyan and orange, respectively. Exosite residues K532 and K533 are colored in yellow and purple, respectively. A) C1; B) C2; C) C3.

**Figure 2 -Figure Supplement 2.**
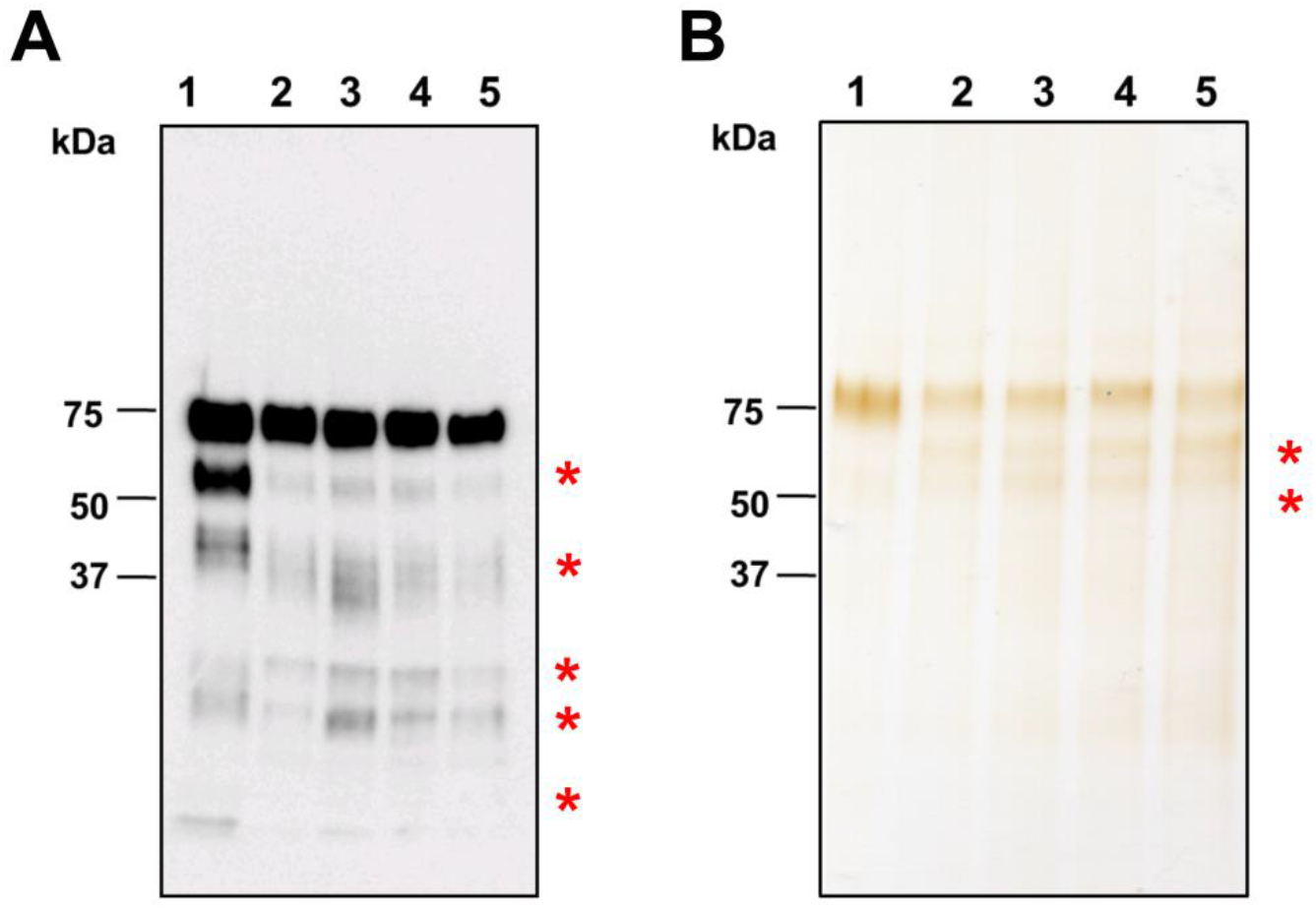
Expression of ADAMTS-5 variants. ADAMTS-5 MDTCS and its Dis domain variants were expressed in HEK293T cells. After 3 days, expression and secretion were analyzed by western blot analysis of conditioned media using an anti-FLAG antibody (Cat. number F1804, Sigma Aldrich). Silver stain of affinity-purified ADAMTS-5 MDTCS and its Dis domain variants. Lanes: 1) wild-type ADAMTS-5 MDTCS; 2) K532A/K533A; 3) K533A; 4) K533H; 5) K532Q/K533Q. Red stars indicate C-terminal processed forms.

**Figure 2 -Figure supplement 3.**
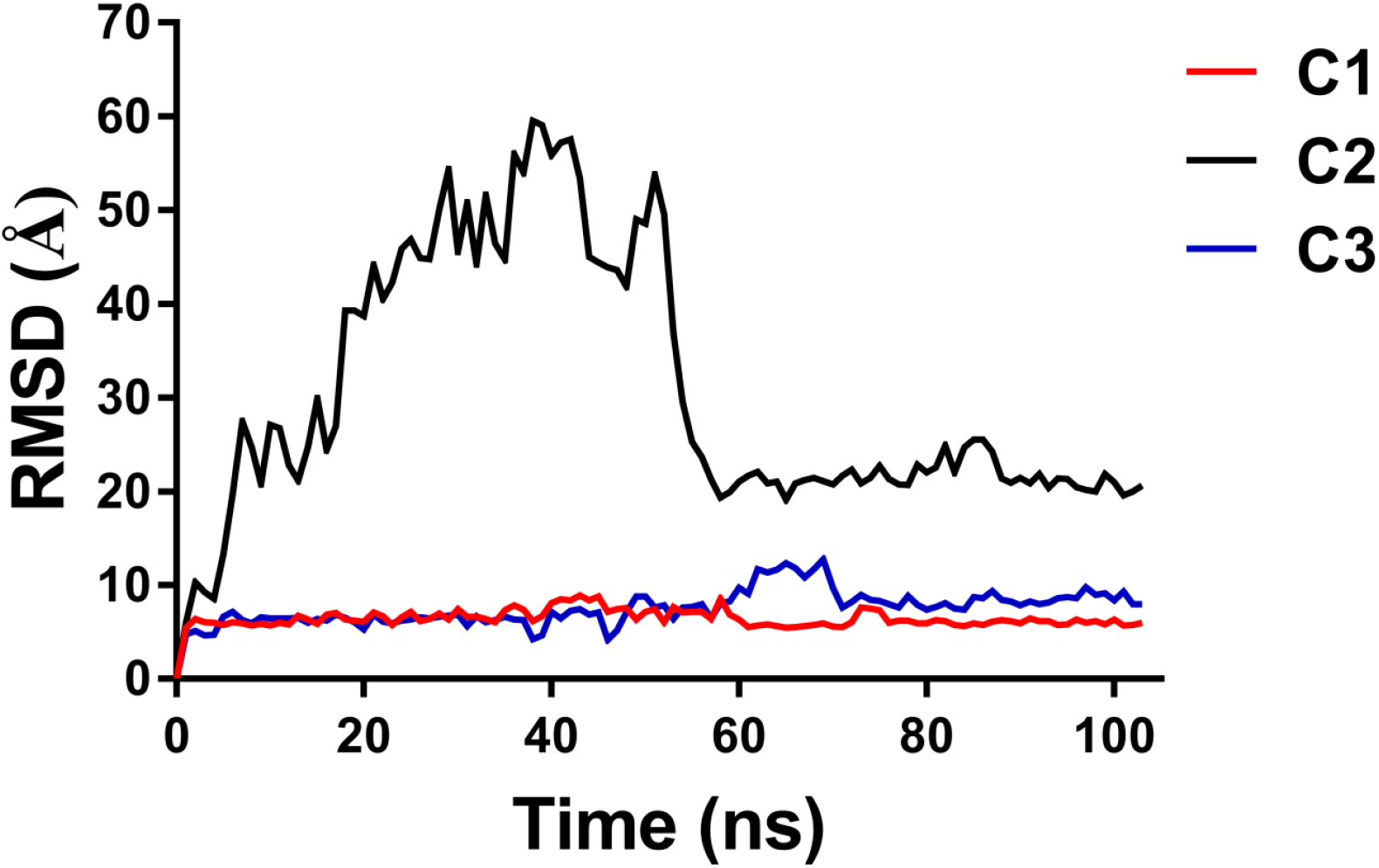
MD analysis for compound 4b in poses C1-C3. RMSD analysis of the heavy atoms of compound 4b in poses C1-C3 during the MD simulation.

**Figure 2 -Table supplement 1.** MM-PBSA results for the three potential orientations of compound 4b into ADAMTS-5. ΔPBSA is the total amount of the electrostatic (EEL), van der Waals (VDW), polar (EPB) and non-polar (PBSUR) solvation free energy. Data are expressed in kcal/mol.

|  | MM-PBSA Method |  |  |  |  |
| --- | --- | --- | --- | --- | --- |
| | VDW | EEL | PBSUR | EPB | ΔPBSA |
| **C1** | -33.8 | -23.8 | -3.9 | 46.7 | -14.7 |
| **C2** | -10.8 | -5.5 | -1.3 | 12.2 | -5.4 |
| **C3** | -30.3 | -32.5 | -3.8 | 59.6 | -6.9 |

